# Bacterial sensing via Neuronal Receptor Initiates Gut Mitochondrial Surveillance for Host Adaptation

**DOI:** 10.1101/2024.10.08.615749

**Authors:** Huimin Liu, Panpan Chen, Xubo Yang, FanRui Hao, Guojing Tian, Zhao Shan, Bin Qi

**Affiliations:** Southwest United Graduate School, Yunnan Key Laboratory of Cell Metabolism and Diseases, Center for Life Sciences, School of Life Sciences, State Key Laboratory of Conservation and Utilization of Bio-resources in Yunnan, Yunnan University, Kunming, China

**Keywords:** Pathogen-like-bacteria, Mitochondrial Surveillance, UPR^mt^, Cell-non-autonomous, Peptidoglycan, Pathogen, Infection, *C. elegans*

## Abstract

Animals exist within a microbial world and are constantly challenged by pathogen infections. Microbe-mediated protection for against infection is the survival strategy for host. However, elucidating specific microbial molecules and understanding how they interact with the host’s intracellular surveillance system for protection is difficult but highly desirable. Here, by establishing “pathogen-like-bacteria” screening system, we identified *E. coli* mutants, including Δ*ymcB*, that act as “pathogen-like-bacteria” to defend animals against *Pseudomonas aeruginosa* PA14 infection by activating UPR^mt^. Additionally, through genetic screening, we identified neuronal transmembrane protein, MDSS-1, that is crucial for sensing Δ*ymcB* and activating intestinal UPR^mt^. Moreover, we demonstrated that MDSS-1 functions as a receptor in ASE neurons, responsible for detecting Δ*ymcB*. It then communicates microbial signals through neuropeptides, GPCR, Wnt signaling and endopeptidase inhibitors to trigger intestinal UPR^mt^, that defends the host animals against infections. Furthermore, Constitutionally activation of MDSS-1 in ASE neurons is sufficient to trigger intestinal UPR^mt^ in animals, resulting in protection against infection. Our study uncovers an intriguing mechanism involving intracellular mitochondrial surveillance, where neuron-intestine crosstalk originates from ASE neurons to detect bacteria and combat pathogens. This study identifies a bacteria-sensing mechanism in neurons that regulates intestinal mitochondrial surveillance pathway for host adaptation.

**Highlights:**

- Establishment of “pathogen-like-bacteria” screening system in *C. elegans*
- Δ*ymcB* promotes animal defend against infections via triggering UPR^mt^
- Neuronal MDSS-1, a single transmembrane protein, detects “pathogen-like-bacteria”
- Activated-MDSS-1 induces distant UPR^mt^ via inter-tissue communication factors

## Introduction

Pathogenic bacterial infections present a significant threat to public health. Animals live in symbiosis with numerous microbe species. The host–microbes’ coevolution are the pivotal events to maintain homeostasis of microbial community (Groussin et al., 2020), which plays an instrumental role in maintaining physiological homeostasis for the health of the host by influencing immunity, metabolism, development and lifespan. Dysbiosis of microbiota is associated with many diseases (Carding et al., 2015) and increased pathogen susceptibility (Pham and Lawley, 2014). Microbe-mediated protection against infection is the survival strategy of the many host species, achieved by preventing pathogen establishment/reproduction or enhancing host immune response (Stevens et al., 2021). Identify specific microbial species or factors in against pathogen infection, and thereby developing genetically engineered probiotics using synthetic biology approaches is a new therapeutic paradigm to benefit host health. However, characterizing individual bacterial species and their mechanisms in hosts are still challenging. Therefore, establishing effective animal models and high-throughput screening approaches to explore the protective bacterial species and the mechanisms by which animals sense these species are highly desirable. Recent studies have demonstrated that the nematode *C. elegans* is excellent system to screen benefit bacteria and study beneficial function of bacteria on animal physiology (Geng et al., 2022; Govindan et al., 2015; Han et al., 2017; He et al., 2023; Lin and Wang, 2017; Sonowal et al., 2017; Tian and Han, 2022; Zhang et al., 2019) and pathogen infection protection (Kissoyan et al., 2019; Kissoyan et al., 2022; Sang et al., 2022; Tsuru et al., 2021).

C. *elegans* reside in environments that are rich in microbes (Dirksen et al., 2020; Dirksen et al., 2016; Felix and Braendle, 2010). Some species of microbes have protection roles for animals (Kissoyan et al., 2019; Kissoyan et al., 2022; Sang et al., 2022; Tsuru et al., 2021), that facilitate adaptation of animals in nature. Therefore, distinguishing bacteria is very critical for animals to maintain physiology homeostasis. Evolutionary evidence strongly suggests that mitochondria descend from bacteria that entered other cell types and became incorporated into their structure (Gogoi et al., 2022). Mitochondria play essential roles in many aspects of the cell’s metabolism, which was also targeted by the microbial toxin or metabolites (Jiang et al., 2012) to affect host physiology, including infection and life-span, via mediating mitochondrial stress (Han et al., 2017; Tian and Han, 2022; Zhou et al., 2021). Beside activating UPR^mt^ by mitochondrial perturbation via knocking down mitochondrial genes (*cco-1, spg-7, atp-2*) (Berendzen et al., 2016; Shao et al., 2016), UPR^mt^ was also triggered by bacteria infection (Pellegrino et al., 2014). However, mitochondrial perturbation upregulates genes that partially overlap with those under pathogen infection, such as genes of innate immunity (Nargund et al., 2012; Pellegrino et al., 2014), indicating that *C. elegans* adopts UPR^mt^ as a surveillance and defense mechanism against pathogenic microbes. Mild mitochondrial stress has the potential to improve host resistance to infection through the activation of mitochondrial stress responses that initiate a protective innate immune response (Campos et al., 2021; Chen et al., 2021). “Pathogen-like-bacteria” that cannot infect host by themselves and activate the host’s UPR^mt^ survival pathway should be ideal species for treating infection disease. Therefore, developing a screening strategy in *C. elegans* to find “pathogen-like-bacteria” and study how animals sense these bacteria is critical for preventing pathogen infection and understanding the adaptation of the host animals in nature.

The protective UPR^mt^ in animals for improving longevity or enhancing infection was reported by disturbing mitochondrion function (mitochondrial perturbations) through artificial methods, such as knocking out of mitochondrial genes (*cco-1, spg-7, atp-2*) in neuron (Berendzen et al., 2016; Shao et al., 2016), neuronal expression polyQ40 or Wnt/EGL-20 (Berendzen et al., 2016; Liu et al., 2022), and perturbing mitochondrial fusion in neurons (Chen et al., 2021). Recent studies using these ideal artificial systems to study tissue-tissue crosstalk have demonstrated that neuronal control of systemic UPR^mt^ activation is critical for coordinating organismal mitochondrial homeostasis and metabolic states to improve animals’ fitness (Berendzen et al., 2016; Chen et al., 2021; Liu et al., 2022; Shao et al., 2016). However, how neuron and their factors sense native bacteria as “pathogen-like-bacteria” to protect animals by inducing intestinal UPR^mt^ is still unclear. Therefore, dissecting the mechanism of animals sensing “pathogen-like-bacteria” to defend against infection via UPR^mt^ is essential to understand the ability to produce genetic diversity in the face of microbial challenge.

In this study, we employ a high-throughput screening system to identify “pathogen-like-bacteria” against infection from an *E. coli* mutant collection. We discovered that several *E. coli* mutants, including Δ*ymcB,* can protect animals against pathogens by activating UPR^mt^. Moreover, we performed a genetic screen to identify genes required for sensing Δ*ymcB* via intestinal UPR^mt^ activation. We identified neuronal transmembrane protein, MDSS-1, that is crucial for sensing Δ*ymcB* and activating intestinal UPR^mt^ to protect against infections. Furthermore, we observed that constitutionally activation of MDSS-1 in ASE neurons is sufficient to trigger intestinal UPR^mt^ in animals, resulting in protection against infection. Notably, ablation of ASE neurons attenuates the intestinal UPR^mt^ activation in response to Δ*ymcB*. Moreover, we found that activation of UPR^mt^ by MDSS-1 relies on canonical UPR^mt^ pathway, neuropeptides, GPCR and Wnt signaling. Additionally, we identified that inhibition of endopeptidase inhibitor activity acts as downstream signaling of MDSS-1 in sensing Δ*ymcB* for inducing UPR^mt^. Overall, we discovered that a neuronal receptor sensing “pathogen-like-bacteria” coordinates the intestinal mitochondrial stress response for pathogen defense, thereby facilitating the adaptation of animals in nature for survival.

## Results

### Establishment of “pathogen-like-bacteria” screening strategy in *C. elegans* using an *E. coli* mutant library

Animals live in symbiosis with numerous microbe species. Nematodes have a characteristic microbiome, including bacterial-feeding nematodes, which represent approximately 80% of all the multicellular animals on earth (Eisenhauer and Guerra, 2019). Therefore, one explanation for their success is the co-evolution of host and symbiotic gut microbiome which benefit animals’ survival, especially under pathogen infection. Certain bacterial species may provide protection to animals against pathogen infection, which is the evolutionary ideal survival strategy for nematodes (Kissoyan et al., 2019; Kissoyan et al., 2022; Sang et al., 2022; Tsuru et al., 2021). Here, we defined these bacteria as "pathogen-like-bacteria" that cannot infect hosts by themselves but can "immunize" animals against pathogen bacteria by activating the host immune pathway and then protect them from infection (Figure 1A and S1A). However, finding specific bacterial species and factors involved in host protection for infection is challenging and highly desirable. Developing genetically engineered probiotics is a promising new therapeutic approach to promote host health (O’Toole et al., 2017). *C. elegans* feed on *E. coli* as their food source in the laboratory. We hypothesized that “pathogen-like-bacteria” could be engineered by mutating genes in *E. coli* that enhance animals’ defense against infection. Therefore, developing a “pathogen-like-bacteria” screening strategy in *C. elegans* by using *E. coli* mutant library (Figure S1B) is an excellent system to find probiotics and study how animals sense probiotics for against infection. Animals were first trained with *E. coli* mutant and then infected with *Pseudomonas aeruginosa* PA14 to assess their survival (Figure S1B). The candidate *E. coli* mutant could protect animals against PA14 infection.

**Figure 1.**
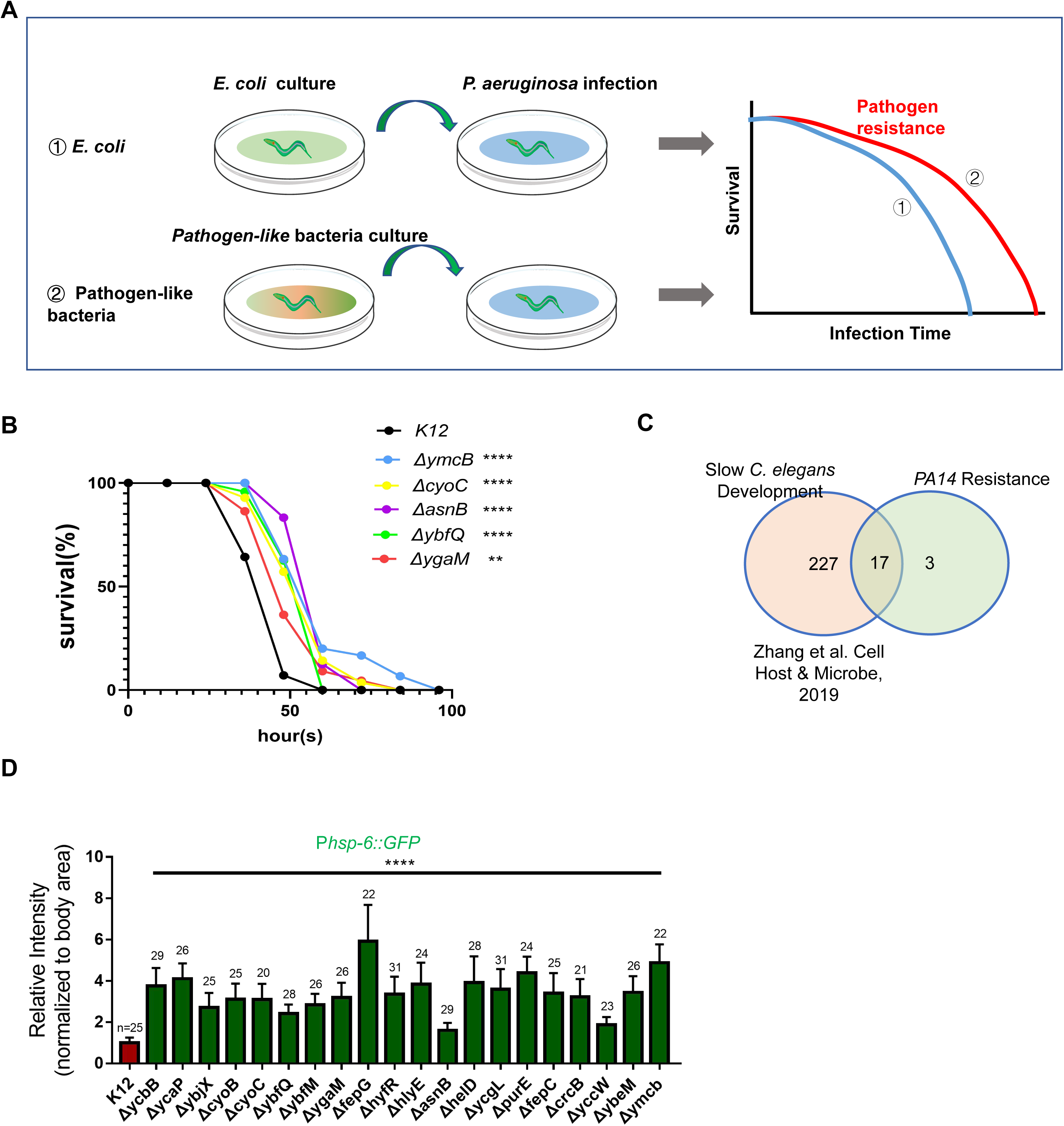
Screen for "pathogen-like-bacteria" that protect animals against infection. **(A)** Cartoon illustration of the established pathogen-like-bacteria pre-training research system in *C. elegans*. The survival rate of animals pre-conditioned with pathogen-like-bacteria and subsequently exposed to *Pseudomonas aeruginosa* PA14 is expected to be higher compared to those exposed to control *E. coli* K12. **(B)** Survival analysis of worms pre-conditioned with *E. coli* K12 or mutant bacteria on PA14. Statistics are in Table S1. **(C)** Diagram showing that animal’s growth is slow with feeding 17 out of 20 *E. coli* mutants obtained by screening. The slow-growth phenotype is derived from the research findings of Zhang et al (Zhang et al., 2019). **(D)** Quantification of P*hsp-6::GFP* reporter expression in animals grown on *E. coli* K12 or pathogen-like-bacteria. For all panels, *p<0.05, **p<0.01, ***p<0.001, ****p<0.0001. n is the number of worms scored. Error bars, ± s.d. All experiments were performed independently at least three times.

### *E. coli* mutant as “pathogen-like-bacteria” protects animals against PA infection by activating UPR^mt^

After screening an *E. coli* single-gene knockout library (Keio collection) (Baba et al., 2006), we discovered that pre-training animals with 20 *E. coli* mutants resulted in increased resistance to PA14 infections, indicating these mutants act as "pathogen-like-bacteria" that protect against infection (Figure 1B and Figure S1C). In a previous study (Zhang et al., 2019), it was discovered that approximately 90% of the *E. coli* mutants, which exhibited delayed development, triggered UPR^mt^ in *C. elegans*. We further explored this aspect and identified that 17 out of the 20 candidates we screened for our research were part of the same bacterial mutant collection (Figure 1C), suggesting that these 20 *E. coli* mutants may also induce UPR^mt^ in our study.

Furthermore, we also confirmed that all 20 *E. coli* mutants activate UPR^mt^ in P*hsp-6::GFP* transgene animals (Figure 1D) which was used to monitor UPR^mt^. Previous reports have demonstrated that UPR^mt^ activation enhances innate immunity against infection (Campos et al., 2021; Pellegrino et al., 2014). Therefore, the UPR^mt^ activation observed in animals trained with the 20 *E. coli* mutants suggests that these bacteria act as "pathogen-like-bacteria" and affect animals’ mitochondrial defense function, ultimately protecting against pathogen infection.

### Animals sense pathogen-like-bacteria **Δ***ymcB* through ATFS-1-dependent UPR^mt^

The simple animal *C. elegans* have developed neuronal sensing system to respond to changing microbial cues and avoid the environmental pathogens (Meisel et al., 2014; O’Donnell et al., 2020; Rhoades et al., 2019; Wu et al., 2023), which is essential for its survival. To investigate whether worms have an aversion response to pathogen-like-bacteria, we performed a food choice assay and found that they prefer wild-type *E. coli* over pathogen-like-bacteria (Figure S2A), indicating that animals have a sensing mechanism to distinguish pathogen-like-bacteria.

To further investigate this sensing mechanism, we simplified the research system by focusing on Δ*ymcB* for two reasons. Firstly, the Δ*ymcB* mutant showed a significant activation of UPR^mt^ in animals (Figure 1D), helping them defend against PA14 infection (Figure 1B). Secondly, previous research has demonstrated that *ymcB* (other name: GfcC) in *Escherichia coli* is crucial for the assembly of the group 4 polysaccharide capsule (Sathiyamoorthy et al., 2011). This capsule is responsible for decorating the surface of bacterial cells and providing structural integrity and protection. Therefore, it is likely that the mutation of *ymcB* disrupts components of the bacterial cell wall, which could be detected by animals.

To confirm whether animals’ response to pathogen-like-bacteria Δ*ymcB*, we first performed food choice assay and found that animals preferred *K12* (Figure 2A). Secondly, we examined the levels of several commonly used stress reporters in animals with Δ*ymcB* feeding and found that only UPR^mt^ (P*hsp-6::GFP*) was strongly induced (Figure 2B-D), while UPR^ER^ (P*hsp-4::GFP*) or immune response (P*irg-5::GFP,* P*sysm-1::GFP)* was not (Figure S2B). These results suggest that animals primarily sense pathogen-like-bacteria Δ*ymcB* through activation of UPR^mt^.

**Figure 2.**
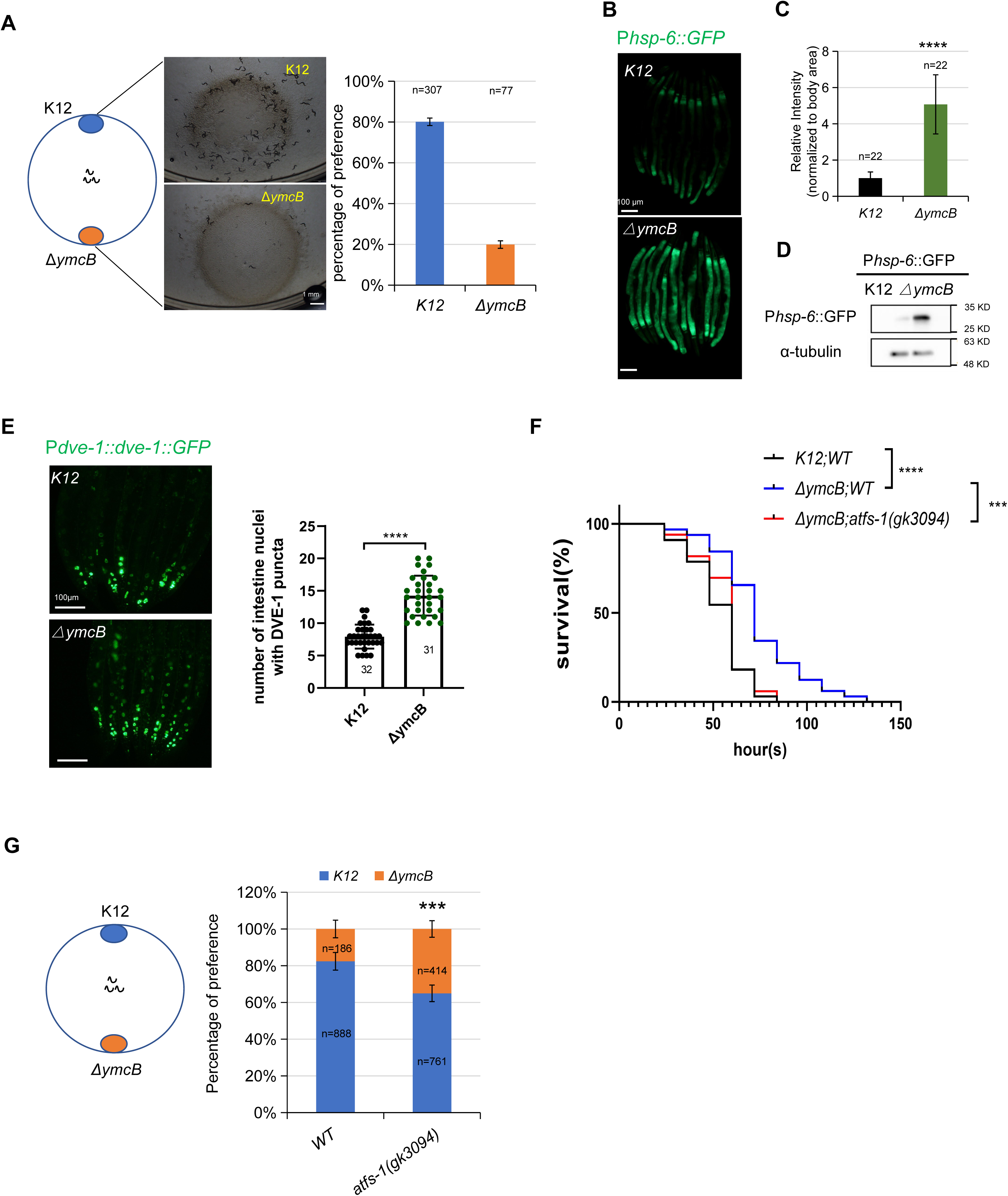
Animals sense pathogen-like-bacteria Δ*ymcB* through ATFS-1-dependent UPR^mt^. **(A)** Microscope images and quantification of selection for *E.coli* K12 or Δ*ymcB* of animals seeded into the middle of two bacteria for 60 hours at 20℃. **(B-D)** Microscope images and fluorescence quantification of P*hsp-6::GFP* reporter expression in animals grown on *E. coli* K12 or Δ*ymcB*. Immunoblot analysis of animals grown on *E. coli K12* or Δ*ymcB* using antibodies against GFP. Anti-GFP detects *Phsp-6::GFP* expression. **(E)** Microscope images and quantification of numbers of intestinal nuclei with P*dve-1::dve-1::GFP* puncta in animals grown on the *E. coli* K12 or Δ*ymcB*. **(F)** Survival analysis of wild-type or *atfs-1(gk3094)* animals pre-conditioned with *E. coli* K12 or Δ*ymcB* on PA14. Statistics are in Table S1. **(G)** Quantification of selection for *E.coli* K12 or Δ*ymcB* of wild-type and *atfs-1(gk3094)* animals seeded into the middle of the two bacteria for 60 hours at 20℃. For all panels, *p<0.05, **p<0.01, ***p<0.001, ****p<0.0001. n is the number of worms scored. Error bars, ± s.d. All experiments were performed independently at least three times.

UPR^mt^ induction requires the key transcription factor ATFS-1, transcriptional co-regulators DVE-1 and UBL-5 (Haynes et al., 2007; Nargund et al., 2012; Tian et al., 2016). RNAi knockdown of *dve-1, atfs-1,* or *ubl-5* significantly suppressed UPR^mt^ in the animals fed with Δ*ymcB* (Figure S2C). Moreover, nuclear localization of P*dve-1::dve-1::*GFP expression was increased in the intestinal cells of animals treated with Δ*ymcB* (Figure 2E). These data indicate that activation of UPR^mt^ in animals with Δ*ymcB* feeding is triggered through the canonical UPR^mt^ pathway.

Next, we test whether ATFS-1-dependent UPR^mt^ activation is essential for animals to sense Δ*ymcB* for their protection. Firstly, we found that Δ*ymcB-*induced protection against PA14 infection was impaired in *atfs-1(gk3094)* mutant (Figure 2F). Secondly, performance for choosing K12 was also decreased in *atfs-1(gk3094)* mutant (Figure 2G), but worms still exhibited some attraction to Δ*ymcB* mutants, although to a lesser extent, implying that other factors may also contribute to the response to Δ*ymcB*. Together, our findings indicate that 1) worms sense pathogen-like-bacteria Δ*ymcB* through ATFS-1-dependent UPR^mt^ for protection against pathogen infection; 2) UPR^mt^ reporter activation may serve as the read-out in animals to dissect the mechanism for sensing Δ*ymcB*.

### Transmembrane protein, MDSS-1, is required for sensing pathogen-like-bacteria

In order to understand how animals’ sense and respond to “pathogen-like-bacteria”, we performed an ethyl methanesulfonate (EMS) mutagenesis screen to identify genes that are required for the UPR^mt^ activation in response to pathogen-like-bacteria treatment. We hypothesized that if key sensing genes were mutated, then the Δ*ymcB* induced protection and behavior would be impaired in mutant animals (Figure S3A). We initiated our EMS screening process by examining approximately 9000-12000 F1 animals harboring the *Phsp-6::GFP* construct, and screened mutant animals in F2 generation which *Phsp-6::GFP* cannot be induced by Δ*ymcB E. coli*. Through this screening, we successfully identified four mutant strains. One of these mutant alleles, *ylf10,* exhibited strong UPR^mt^ suppression with Δ*ymcB* feeding (Figure 3A-B). Whole-genome sequencing and sibling subtraction method identified a missense mutation at the 91^th^ amino acid (GGA → GAA[Gly91Glu]) of the gene *C32C4.3* in *ylf10* mutant, which encodes an uncharacterized single transmembrane protein (Figures 3A, Figure 4A). As C32C4.3 is an uncharacterized single transmembrane protein involved in “pathogen-like-bacteria” response and UPR^mt^ regulation, therefore, we named C32C4.3 as <u>m</u>itochon<u>d</u>rial <u>s</u>urveillance <u>s</u>ensor gene in sensing bacteria (*mdss-1*). We then generated a translation-terminated mutant (namely, *ylf14*) that the stop codon is immediately following the start codon of C32C4.3 by using CRISPR-Cas9-mediated gene editing (Figures 3A). Next, we confirmed that 1) UPR^mt^ induction in animals feeding Δ*ymcB* is suppressed in *mdss-1* mutant (*ylf10* and *ylf14*) (Figures 3B-C) or through *mdss-1* RNAi (Figure S3B), 2) UPR^mt^ suppression in *mdss-1* mutation was rescued in transgenic animals expressing *mdss-1* by using either its own promoter (Figure 3D) or a ubiquitously expressed *rpl-28* promoter (Figure S3C), indicating that *mdss-1* is required for the Δ*ymcB* induced UPR^mt^. UPR^mt^ decreases by approximately 2-fold when *mdss-1* is mutated (Figure 3C); however, expression of the *Pmdss-1::mdss-1* transgene induces only a 1.3-fold increase (Figure 3D). This suggested that *Pmdss-1::mdss-1* transgene partially rescues the mutant phenotype.

**Figure 3.**
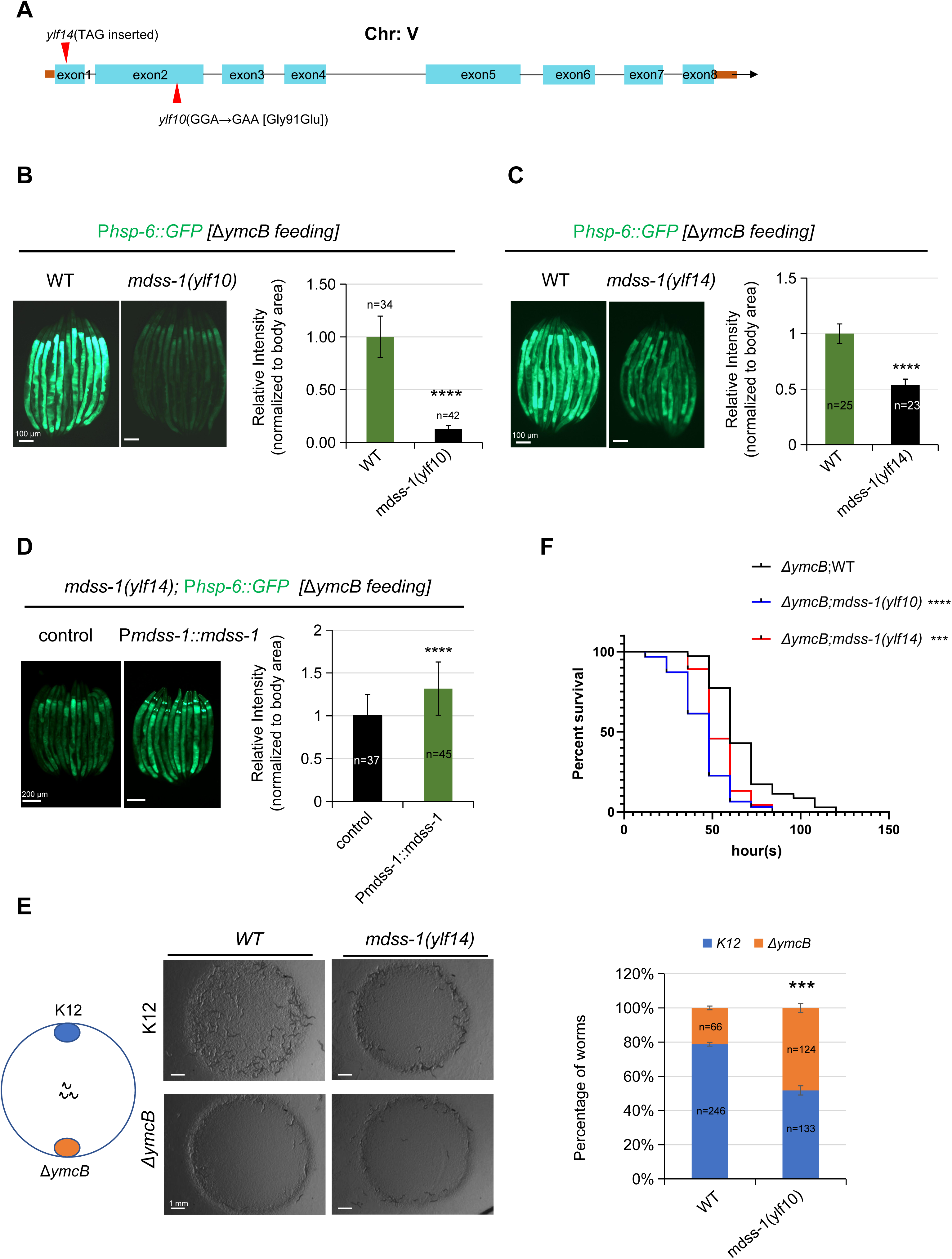
Transmembrane protein MDSS-1 is required for sensing pathogen-like-bacteria. **(A)** A schematic of the two mutant alleles *ylf10* or *ylf14* of *mdss-1* in *C. elegans* genome. **(B)** Microscope images and fluorescence quantification of P*hsp-6::GFP* reporter expression in wild-type or *mdss-1(ylf10)* animals grown on the *E. coli* K12 or Δ*ymcB*. **(C)** Microscope images and fluorescence quantification of P*hsp-6::GFP* reporter expression in wild-type or *mdss-1(ylf14)* animals grown on *E. coli K12* and Δ*ymcB*. **(D)** Microscope images and fluorescence quantification of P*hsp-6::GFP* reporter expression in *mdss-1(ylf14)* animals with or without P*mdss-1::mdss-1::mcherry* transgene grown on Δ*ymcB*. Animals carrying transgenes discriminated by P*odr-1::GFP*. **(E)** Microscope images and quantification of selection for *E. coli K12* or Δ*ymcB* of wild-type and *mdss-1(ylf14)* animals seeded into the middle of the two bacteria for 60 hours at 20℃. **(F)** Survival analysis of wild-type and *mdss-1* mutant animals pre-conditioned with Δ*ymcB* on PA14. Statistics are in Table S1. For all panels, *p<0.05, **p<0.01, ***p<0.001, ****p<0.0001, n is the number of worms scored. Error bars, ± s.d. All experiments were performed independently at least three times.

**Figure 4.**
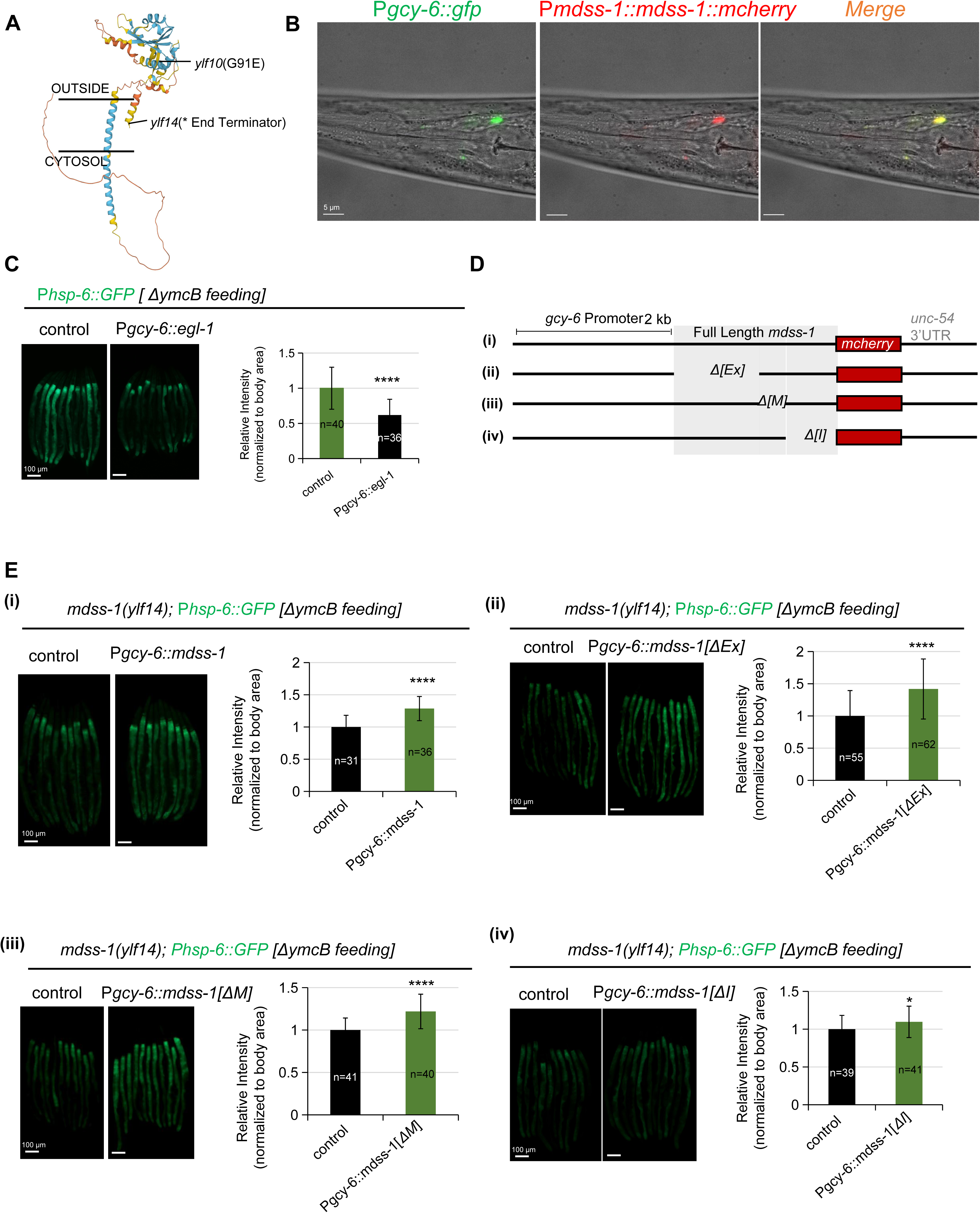
The intracellular region of MDSS-1 in ASE neurons plays a crucial role in facilitating the nonautonomous activation of UPRmt in response to Δ*ymcB*. **(A)** A schematic of MDSS-1 transmembrane protein. The line indicates the amino acid sites corresponding to the two mutations in *ylf10* and *ylf14*. **(B)** Microscope images of P*gcy-6::GFP* and P*mdss-1::mdss-1::mcherry* in wild-type animals. The P*gcy-6::GFP* transgene shows specific expression in ASE neurons. **(C)** Microscope images and fluorescence quantification of P*hsp-6::GFP* reporter expression in animals with or without P*gcy-6::egl-1::mkate2* transgene grown on Δ*ymcB*. **(D)** A schematic of *mdss-1* isoforms with varying lengths driven by *gcy-6* promoter, including i) full length *mdss-1*, ii) Δ*[Ex]* (extracellular segment deletion) *mdss-1*, iii)Δ*[M]*(transmembrane segment deletion) *mdss-1*, iv) Δ*[I]*(intracellular segment deletion) *mdss-1*. **(E)** Microscope images and fluorescence quantification of P*hsp-6::GFP* reporter expression in *mdss-1(ylf14)* animals with or without *mdss-1* isoforms transgen as described in (D) grown on Δ*ymcB*. For all panels, *p<0.05, **p<0.01, ***p<0.001, ****p<0.0001, n is the number of worms scored. Error bars, ± s.d. All experiments were performed independently at least three times.

To further investigate *mdss-1*’s role in sensing “pathogen-like-bacteria”. We conducted a food choice assay, which show that *mdss-1*(*ylf14*) mutant performance for choice K12 was decreased (Figure 3E). The survival assay demonstrated that Δ*ymcB-*induced protection in animals against PA14 infection is impaired in *mdss-1(ylf14)* mutant (Figure 3F). Moreover, *mdss-1(ylf14)* also exhibited significant suppression of UPR^mt^ when fed with other "pathogen-like bacteria" in addition to Δ*ymcB* (Figure S3D). Together, our data indicates that *mdss-1* plays a crucial role in sensing pathogen-like-bacteria and triggering UPR^mt^ to resist pathogen infection.

The previous study demonstrated that neuronal expression of polyQ40 results in impaired mitochondrial function and the initiation of a non-cell-autonomous UPR^mt^ (Berendzen et al., 2016). We found that the activation of intestinal UPR^mt^, induced by polyQ40 expression in neurons, remained unaffected by *mdss-1* mutations (Figure S3E). This result indicates that MDSS-1 primarily function as a sensor for “pathogen-like-bacteria” rather than responding to general cellular metabolic or mitochondrial stress.

### MDSS-1 in ASE neurons plays a crucial role in response to pathogen-like-bacteria

Animals have the ability to respond to bacteria by generating adaptive behaviors, which are regulated by neuronal homeostasis and downstream effects (Meisel et al., 2014; O’Donnell et al., 2020; Rhoades et al., 2019; Wu et al., 2023). Neurons also play a critical role in regulating cell-non-autonomous UPR^mt^ for stress response (Berendzen et al., 2016; Chen et al., 2021; Frakes et al., 2020; Liu et al., 2022). However, the mechanism by which the neurons sense bacteria and signal to the distal tissues to regulate animals’ physiology remains unclear.

To further investigate how MDSS-1 sense pathogen-like-bacteria, we utilized AlphaFold (https://alphafold.com) to predict its protein structure, which revealed that MDSS-1 is a single transmembrane protein with a receptor L domain located in the intracellular region (Figure 4A). We also searched for the expression level of C32C4.3 in the CeNGEN database (Hammarlund et al., 2018), and found that it is highly expressed in ASE neurons. Subsequently we examined the expression pattern of *mdss-1* by constructing transgenic animals, and confirmed that it is specifically expressed in ASE neurons through co-localization analyses of P*mdss-1::mdss-1::mcherry* reporter with ASE neuronal marker P*gcy-6::GFP* (Figure 4B). Therefore, we constructed transgenic animals that over-express *egl-1*, a cell death activator (Conradt and Horvitz, 1998), in ASE neurons, for ablating ASE neurons. We observed that ablation of ASE neurons significantly suppressed the intestinal UPR^mt^ activation in response to pathogen-like-bacteria, Δ*ymcB* (Figure 4C). Additionally, we observed that the suppression of UPR^mt^ in the intestine of *mdss-1*(*ylf14*) mutant worms by Δ*ymcB* feeding was rescued by the expression of *mdss-1* in ASE neurons driven by the *gcy-6* promoter (Figure 4E-i), indicating that MDSS-1 is functional in ASE neurons and can sense pathogen-like-bacteria to stimulate intestinal UPR^mt.^. In summary, these findings collectively suggest that MDSS-1 in ASE neurons plays a crucial role in the activation of intestinal UPR^mt^ in response to pathogen-like-bacteria, Δ*ymcB*.

### The intracellular region of MDSS-1 in ASE neurons plays a crucial role in facilitating the non-autonomous activation of UPR^mt^ in response to **Δ***ymcB*

In order to investigate the role of different regions of MDSS-1 in the response of animals to pathogen-like bacteria, we generated transgenic animals expressing various isoforms of *mdss-1* under the control of the *gcy-6* promoter. These isoforms included: i) full-length *mdss-1*, ii) *mdss-1* with the extracellular segment deletion (Δ*[Ex]*), iii) *mdss-1* with the transmembrane segment deletion (Δ*[M]*), and iv) *mdss-1* with the intracellular segment deletion (Δ*[In]*) (Figure 4D). We found that the expression of *mdss-1* with the intracellular fragment (Figure 4E i-iii) led to a significant rescue of UPR^mt^ in animals feeding Δ*ymcB*, whereas the efficiency of *mdss-1* lacking the intracellular fragments was less effective (Figure 4E iv). These findings suggest that extracellular and intracellular *mdss-1* play distinct roles in ASE neurons in transmitting signals to activate UPR^mt^ in the intestine. Moreover, it is highly likely that intracellular MDSS-1 facilitates the nonautonomous activation of UPR^mt^ in response to Δ*ymcB*.

To investigate whether Δ*ymcB* affects the expression levels of MDSS-1 and thus influences the activation of distal UPR^mt^, we examined the expression levels of MDSS-1 in the presence of K12 and Δ*ymcB*. We found that no significant difference in MDSS-1 expression levels between the two strains (Figure S4). This suggests that the regulatory mechanism through which MDSS-1 controls UPR^mt^ in response to Δ*ymcB* does not involve alterations in its expression.

In summary, these findings imply that MDSS-1 may act as potential receptors, responsible for sensing Δ*ymcB, and* subsequently transmitting UPR^mt^ signals through the intercellular region of MDSS-1, rather than modulating its expression levels.

### MDSS-1 acts as a potential receptor in UPR^mt^ signal transmission

Single-pass transmembrane proteins play a crucial role in many cellular signaling pathways, including signal transduction, cell adhesion, communications, and immune response (Pahl et al., 2013). We have demonstrated that MDSS-1, a single-pass transmembrane protein, is essential for initiating the activation of UPR^mt^ in response to “pathogen-like-bacteria”. We speculated that MDSS-1 may function as a potential receptor in ASE neurons, facilitating the induction of UPR^mt^ in response to “pathogen-like bacteria”.

If MDSS-1 is a receptor, this would mean that constitutive activation of MDSS-1 should also activate signaling and induce the intestinal UPR^mt^ for protection against infection without pathogen-bacteria training. To test this, we constructed transgenic animals with extrachromosomal arrays expressing *mdss-1* under the control of its own promoter or the ASE neuronal specific expressed promoter (*gcy-6*). We found that overexpression of *mdss-1* in ASE neurons was sufficient to activate intestinal UPR^mt^ even in the absence of pathogen-like-bacteria (Figure 5A-B). Moreover, we observed that the classical UPR^mt^ components ATFS-1, DVE-1, and UBL-5 were all required for the cell nonautonomous UPR^mt^ response (Figure S5A). Furthermore, we found that the activation of UPR^mt^ through the overexpression of *mdss-1* in ASE neurons was sufficient to provide resistance against PA14 infection (Figure 5C-D).

**Figure 5.**
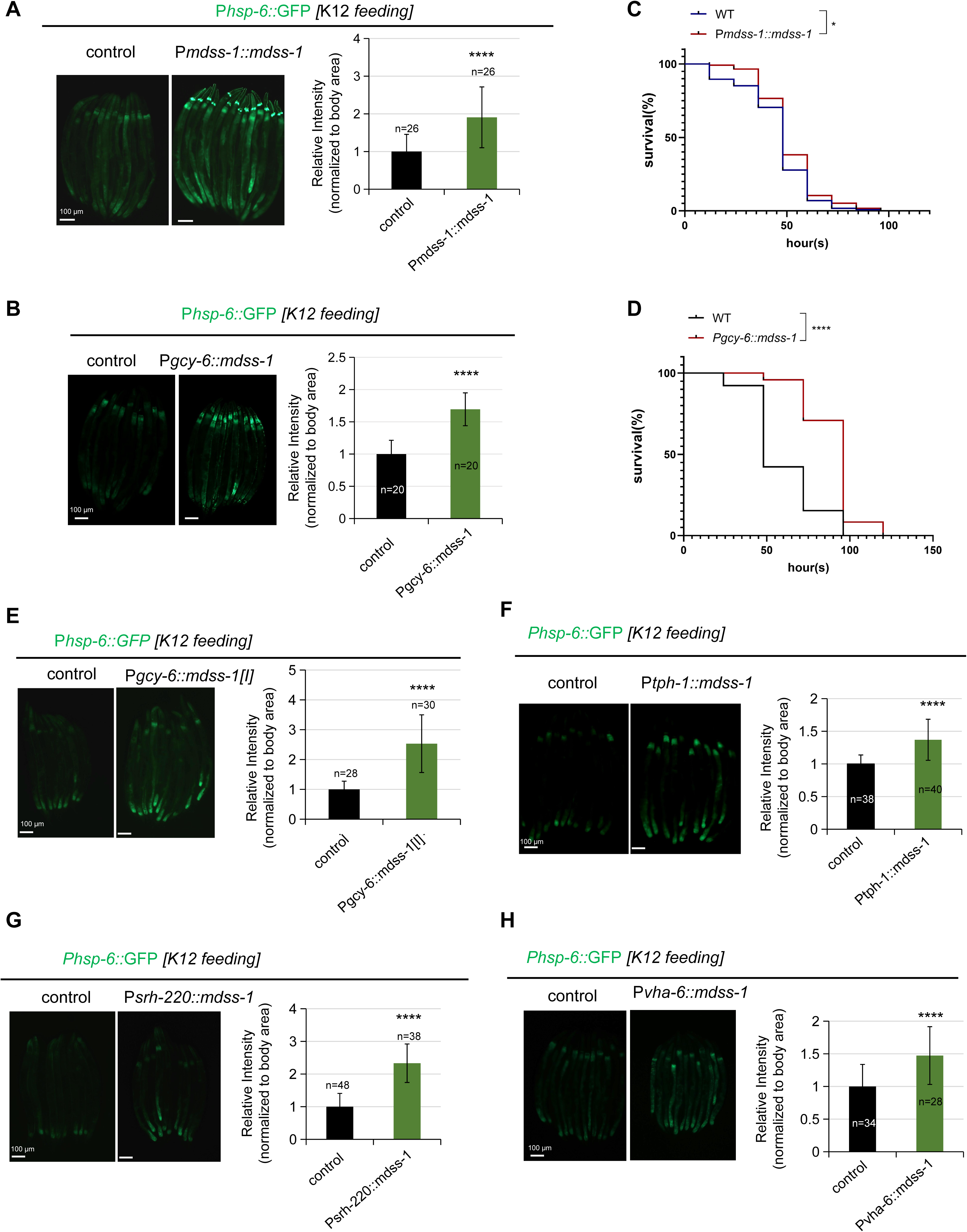
MDSS-1 acts as a potential receptor in UPR^mt^ signal transmission. **(A)** Microscope images and fluorescence quantification of P*hsp-6::GFP* reporter expression in animals with or without P*mdss-1::mdss-1::mcherry* transgene grown on *E. coli K12.* Animals carrying transgenes discriminated by P*odr-1::GFP*. **(B)** Microscope images and fluorescence quantification of P*hsp-6::GFP* reporter expression in animals with or without P*gcy-6::mdss-1::mcherry* t transgene grown on *E. coli K12*. **(C)** Survival analysis of wild-type animals with or without P*mdss-1::mdss-1::mcherry* transgene on PA14. Statistics are in Table S1. **(D)** Survival analysis of wild-type animals with or without P*gcy-6::mdss-1::mcherry* transgene on PA14. Statistics are in Table S1. **(E)** Microscope images and fluorescence quantification of *Phsp-6::GFP* reporter expression in animals with or without P*gcy-6::mdss-1[I]::mcherry* transgene (intracellular *mdss-1*) grown on *E. coli K12*. **(F)** Microscope images and fluorescence quantificatio n of P*hsp-6::GFP* reporter expression in animals with or without P*tph-1::mdss-1::mcherry* transgene grown on *E. coli* K12. **(G)** Microscope image animals and fluorescence quantification of P*hsp-6::GFP* reporter expression in animals with or without P*srh-220::mdss-1::mcherry* transgene grown on *E. coli* K12. **(H)** Microscope images and fluorescence quantification of *Phsp-6::GFP* reporter expression in animals with or without P*vha-6::mdss-1::mcherry* transgene grown on *E. coli* K12. For all panels, *p<0.05, **p<0.01, ***p<0.001, ****p<0.0001, n is the number of worms scored. Error bars, ± s.d. All experiments were performed independently at least three times.

Considering the distinct roles of the extracellular and intracellular regions of MDSS-1 (Figure 4E), we overexpressed either the extracellular or intracellular region of MDSS-1 specifically in ASE neurons. Interestingly, only the intracellular region of MDSS-1 significantly activates UPR^mt^ in the intestine (Figure 5E), whereas extracellular MDSS-1 has no effect under K12 feeding conditions (Figure S5B). This suggests that overexpression of intracellular region of MDSS-1 in ASE neurons triggers the signals to activate intestinal UPR^mt^.

Moreover, if MDSS-1 functions as a receptor in ASE neurons to activate intestinal UPR^mt^, it is likely that the constant activation of MDSS-1 in other tissues would also initiate signaling pathways and result in the induction of intestinal UPR^mt^. Thus, we conducted experiments involving the overexpression of MDSS-1 in the other neurons (ADL neurons and ADF), as well as in intestine. Our results showed that UPR^mt^ was indeed activated in other two neurons (Figure 5F-G) and intestine (Figure 5H). Further analysis revealed that the activation of UPR^mt^ through the expression of *mdss-1* in the intestine involved the participation of transcription factors ATFS-1, DVE-1, and UBL-5 (Figure S5C).

Taken together, these findings support the conclusion that MDSS-1 acts as a potential receptor in sensing “pathogen-like bacteria” for UPR^mt^ activation.

### MDSS-1-mediated UPR^mt^ activation depends on various factors involved in the inter-tissue communication

To gain further insights into the mechanism underlying the activation of intestinal UPR^mt^ by neuronal MDSS-1, we investigated whether this process depends on various factors involved in the inter-tissue communication of mitochondrial stress in *C. elegans*, such as neuropeptides, GPCR, and Wnt signals.

Neuropeptides have been shown to play a crucial role in mediating cell communication, including glial-mediated induction of peripheral UPR^ER^ (Frakes et al., 2020) and the response of distal cells to mitochondrial stress in nerves (Shao et al., 2016). To investigate whether MDSS-1-mediated intestinal UPR^mt^ depends on neuropeptides, we conducted a screening of various neuropeptides (Table S2) based on two criteria: i) enrichment in ASE neurons, and ii) down-regulation in *mdss-1* mutant animals. Through screening, we discovered that the induced UPR^mt^ in animals via overexpressing *mdss-1* is suppressed when *flp-2, flp-5,* or *nlp-3* and their respective receptors are knocked down (Figure S6A-B, Figure 6A). This suggests that the activation of UPR^mt^ by MDSS-1 is dependent on the neuropeptides *flp-2, flp-5,* or *nlp-3*. Furthermore, we observed predominant expression of *nlp-3* in ASE neurons (Figure S6C), and the specific overexpression of *nlp-3* in ASE neurons induced intestinal UPR^mt^ (Figure 6B). Additionally, RNAi knockdown of *nlp-3* suppressed Δ*ymcB*-induced UPR^mt^ (Figure 6C). These results strongly indicate that the activation of UPR^mt^ by MDSS-1 relies on neuropeptides, particularly NLP-3.

**Figure 6.**
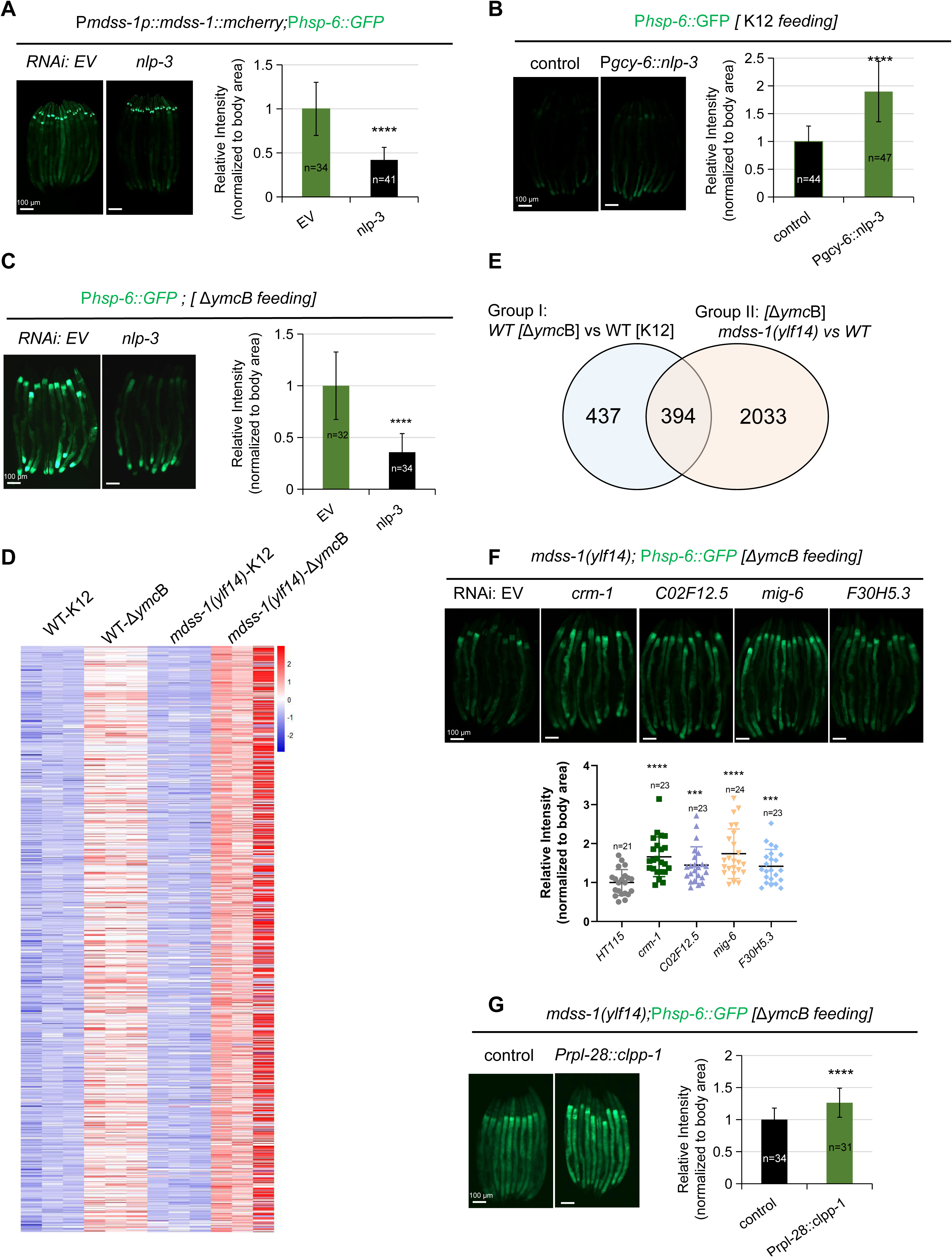
The activation of MDSS-1-mediated UPR^mt^ involves the participation of inter-tissue communication factors and endopeptidase inhibitors. **(A)** Microscope images and fluorescence quantification of P*hsp-6::GFP* reporter expression in animals containing P*mdss-1::mdss-1::mcherry* transgene grown on empty vector (EV) or *nlp-3* (RNAi). Animals carrying transgenes discriminated by P*odr-1::GFP*. **(B)** Microscope images and fluorescence quantification of P*hsp-6::GFP* reporter expression in animals with or without P*gcy-6::nlp-3* transgene grown on *E. coli* K12. **(C)** Microscope images and fluorescence quantification of P*hsp-6::GFP* reporter expression in animals feeding with either empty vector (EV) or *nlp-3* (RNAi) followed by exposure to Δ*ymcB*. **(D)** Heatmap of differentially upregulated genes induced by Δ*ymcB* in group I(wild-type animals grown on K12 transferred to K12 and Δ*ymcB* respectively) and group II(*mdss-1(ylf14)* animals grown on K12 transferred to K12 and Δ*ymcB* respectively). Genes with an adjusted p value< 0.05 were selected as differentially expressed genes. **(E)** Venn diagram of numbers of differentially upregulated genes in wild-type and *mdss-1(ylf14)* animals grown on *E. coli* K12 transferred to Δ*ymcB* for 24 hours. Genes with an adjusted p value< 0.05 were selected as differentially expressed genes. **(F)** Microscope images and fluorescence quantification of P*hsp-6::GFP* reporter expression in *mdss-1(ylf14)* animals grown on either empty vector(EV), *crm-1*(RNAi), *C02F12.5* (RNAi), *mig-6* (RNAi) or *F30H5.3* (RNAi) followed by exposure to Δ*ymcB*. **(G)** Microscope images and fluorescence quantification of P*hsp-6::GFP* reporter expression in *mdss-1(ylf14)* animals with or without P*rpl-28::clpp-1* grown on Δ*ymcB*. For all panels, *p<0.05, **p<0.01, ***p<0.001, ****p<0.0001, n is the number of worms scored. Error bars, ± s.d. All experiments were performed independently at least three times.

GPCR/FSHR-1 (Kim and Sieburth, 2020) and GPCR/SRZ-75 (Liu et al., 2022) in neurons are involved in mediating systemic UPR^mt^ activation in *C. elegans*. Other studies identified that retromer-dependent Wnt/EGL-20 secretion and the Wnt secretion receptor (*mig-14*) are crucial for the cell-non-autonomous activation of UPR^mt^ (Liu et al., 2022; Zhang et al., 2021). To investigate whether MDSS-1-mediated intestinal UPR^mt^ depends on GPCR and Wnt signaling, we conducted experiments to knock down the expression of *fshr-1, srz-75, egl-20* or *mig-14.* We found that induced UPR^mt^ in animals with overexpression of *mdss-1* was significantly inhibited when *fshr-1, srz-75, egl-20* or *mig-14* was knocked down (Figure S6D-E). Moreover, UPR^mt^ in animals with neuronal overexpression of *egl-20* was suppressed by *mdss-1* RNAi knockdown (Figure S6F). This finding strongly suggests that the activation of UPR^mt^ by MDSS-1 depends on GPCR and Wnt signaling, and MDSS-1 has the potential to interact with Wnt/EGL-20 pathwary.

Altogether, our results indicate that the activation of intestinal UPR^mt^ by neuronal MDSS-1 is dependent on various factors involved in the inter-tissue communication of mitochondrial stress, such as neuropeptides, GPCR, and Wnt signals.

### Inhibition of endopeptidase inhibitor activity as downstream signaling of MDSS-1 in sensing **Δ***ymcB* for inducing UPR^mt^

To understand the MDSS-1 regulated downstream effects in sensing pathogen-like-bacteria Δ*ymcB* for intestinal UPR^mt^ activation on a whole-genome level, we performed RNA-seq analysis in wild-type and *mdss-1(ylf14)* mutant animals with or without Δ*ymcB* feeding condition. In group I, we observed that about 800 genes were up-regulated (Figure 6D-E) and 762 genes were down-regulated (Figure S7A-B) in wild-type animals trained with Δ*ymcB* compared with K12 fed animals (Table S3). In group II, about 2400 genes were up-regulated (Figure 6D-E) and 600 genes were down-regulated (Figure S7A-B) in *mdss-1(ylf14)* mutant animals compared with N2 animals feeding Δ*ymcB* (Table S3). About half of Δ*ymcB* induced gene (394/437+394) are overlapped in both groups (Figure 6E), indicating that about half of genes involved in respond to Δ*ymcB* are regulated by *mdss-1*. For all of overlapped genes (394), their expression level is higher in *mdss-1(ylf14)* mutant compared with N2 animals feeding Δ*ymcB* (Figure 6D-E), suggested that those 394 Δ*ymcB* regulated genes are negatively regulated by MDSS-1.

Further GO term enrichment analysis revealed that the majority of the overlapping differentially up-regulated genes were involved in peptidase inhibitor activity (Figure S7C, Table S4), suggesting that endopeptidase inhibitor activity signaling may be downstream of MDSS-1 and may negatively regulate UPR^mt^ activation.

CLPP-1 is a serine-type endopeptidase that degrading the unfolded protein into peptide fragments (Figure S7D) and functions upstream in the UPR^mt^ (Haynes et al., 2007). Our RNA-seq data identified that serine-type endopeptidase inhibitors were enriched in *mdss-1(ylf14)* mutant animals with Δ*ymcB f*eeding (Figure S7C, S7E). This suggests that activity of endopeptidase inhibitors may be increased in *mdss-1(ylf14)* mutants, thereby inhibiting the activity of CLPP-1 and suppressing the UPR^mt^ (Figure S7D). To test this hypothesis, i) we selected several serine-type endopeptidase inhibitors which highly expressed in *mdss-1* mutant (Figure S7E), and found that UPR^mt^ suppression was partially rescued in *mdss-1* mutant by knocking down *crm-1, C02F12.5, mig-6, F30H5.3* (Figure 6F), indicating that these highly expressed endopeptidase inhibitors in *mdss-1(ylf14)* mutant inhibit CLPP-1’s function for activating UPR^mt^. ii) we overexpressed CLPP-1 in the *mdss-1* mutant, and found that UPR^mt^ was induced in the *mdss-1* mutant when CLPP-1 was overexpressed under Δ*ymcB* feeding conditions (Figure 6G). This provides additional evidence to support the notion that MDSS-1 inhibits the expression of endopeptidase inhibitors after sensing Δ*ymcB*, thereby activating CLPP-1’s function to induce UPR^mt^.

Considering that neuronal expression of *mdss-1* activates intestinal UPR^mt^ in the presence of Δ*ymcB*, we hypothesized that neuronal *mdss-1* may inhibit the intestinal expression of endopeptidase inhibitors. To investigate this, we analyzed the expression patterns of these endopeptidase inhibitors (*crm-1, C02F12.5, mig-6, F30H5.3*) through a published database (https://worm.princeton.edu/) (Kaletsky et al., 2018). The results demonstrate that although these inhibitor genes are not specifically expressed in the gut, they show a general enrichment in the intestine compared to other tissues (Figure S7F). This suggests that neuronal *mdss-1* might play a role in modulating the expression of these endopeptidase inhibitors in the intestine, potentially contributing to the activation of UPR^mt^.

In summary, our data strongly suggest that MDSS-1 acts to inhibit the expression of endopeptidase inhibitors following detection of Δ*ymcB*, leading to the activation of CLPP-1 and subsequent induction of UPR^mt^.

## Discussion

It is essential for all animals to respond to the complex microbial world and protect themselves against pathogenic infections. However, identifying specific microbial species and discovering their sensing mechanisms in host that modulate host intracellular surveillance remains a challenge. By establishing “pathogen-like-bacteria” screening system, we found that *E. coli*, when several genes are deleted, absence of peptidoglycan, thereby protecting animals against pathogen infection by activating of UPR^mt^. On the host side, we developed a forward genetics screen system by using UPR^mt^ activation as readout for evaluating the sensing of “pathogen-like-bacteria”, and identified that MDSS-1 as a potential transmembrane receptor that senses “pathogen-like-bacteria” in ASE neuron, which in turn activates intestinal mitochondrial surveillance for pathogen defense. Together, our findings have not only identified specific *E. coli* mutants but have also discovered their sensing mechanisms in neurons that modulate mitochondria homoeostasis in a cell-non-autonomous manner to protect the host against pathogen infections (Figure 7).

**Figure 7.**
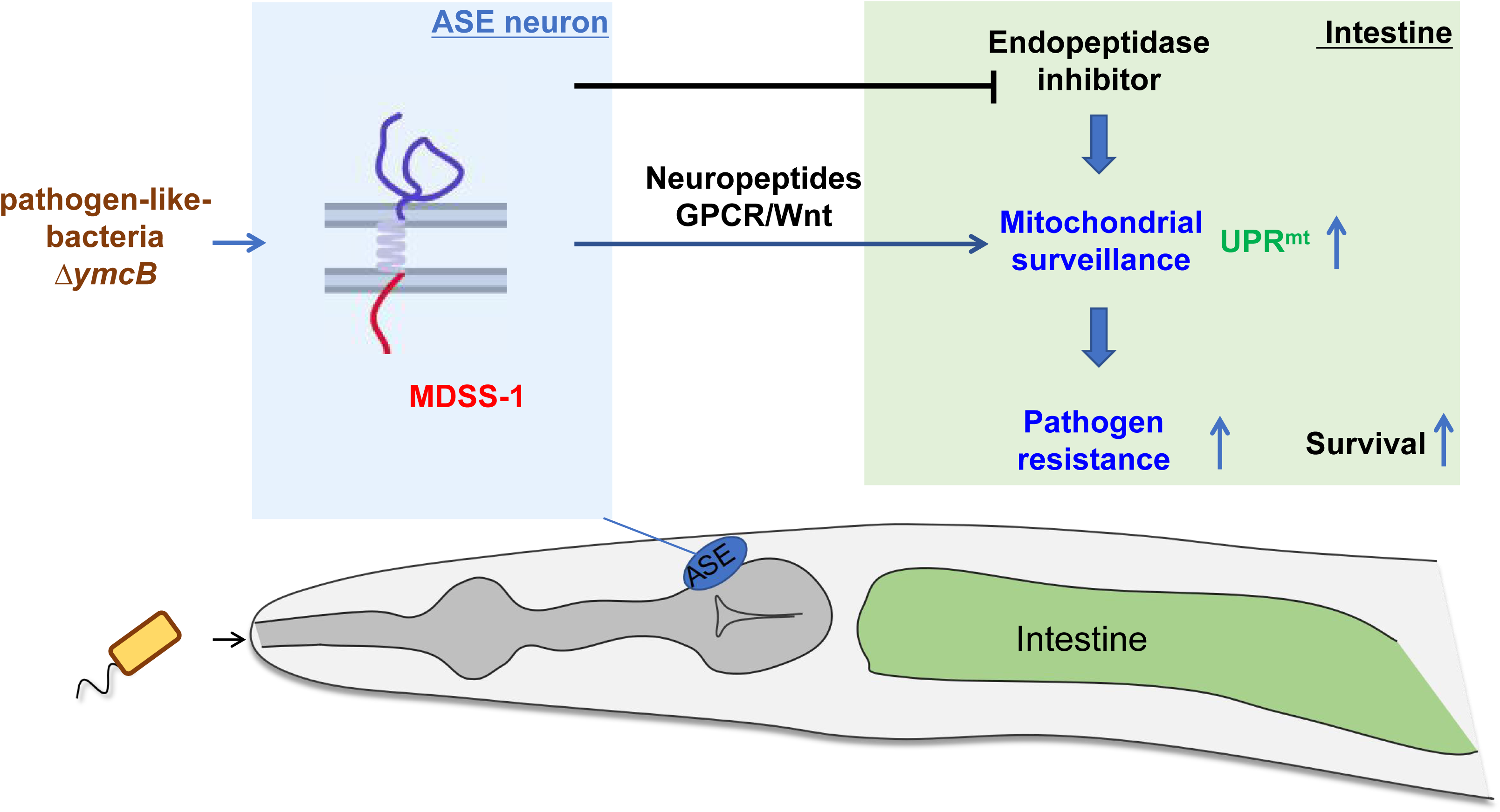
A schematic model illustrates how the neuronal receptor, MDSS-1, senses pathogen-like-bacteria, which triggers intestinal UPR^mt^ for defending against pathogens. Single transmembrane protein, MDSS-1, acts as a potential receptor in ASE neuron to transmit the microbial signals to the intestinal UPR^mt^. This is achieved through inter-tissue communication factors like neuropeptides, GPCR, Wnt signaling, and endopeptidase inhibitors. The induction of intestinal UPR^mt^, triggered by neural MDSS-1 sensing in response to pathogen-like bacteria, helps the animal defend against pathogen infections.

Previous studies have revealed that host microbiota generally play a protective role in preventing or fighting infection (Kissoyan et al., 2019; Kissoyan et al., 2022; Sang et al., 2022; Tsuru et al., 2021). The genetically engineered probiotics is a new therapeutic paradigm to treating infection disease (O’Toole et al., 2017), which will be imperative to ensure that engineered commensals are safe and don’t share their DNA with other bacteria in the environment. *E. coli* belong to enterobacteria is high prevalence in the gut of humans and *C. elegans*. Therefore, one promising strategy to prevent pathogen infection and boost immunity is to engineer *E. coli* strains as "pathogen-like-bacteria" for immunizing and protecting animals (Figure 1A). By establishing “pathogen-like-bacteria” screening system in *C. elegans*, we systematically identified bacterial genes whose deletion leads to pathogen resistance. Remarkably, despite the heterogeneous nature of these bacterial genes, all of these mutants were observed to activate UPR^mt^, suggesting that they may share a common factor which triggers this response. The screening system developed in this study provides a valuable approach for identifying individual bacterial species or engineered commensals that boost host immunity by modulating intracellular stress surveillance pathways to counter infections.

Bacteria-mediated host protection can be mediated in various ways, such as resource competition (Kamada et al., 2012), interference competition (King et al., 2016), or the host immune response (Brown et al., 2013). When infected, host cells can employ intracellular surveillance or stress response programs to detect pathogens (Dunbar et al., 2012; McEwan et al., 2012; Melo and Ruvkun, 2012), and initiate a defense response through innate immunity. However, the mechanisms by which modulation of host intracellular surveillance pathway through sensing these bacteria to better control pathogen infection remain unclear. In our study, we found that an *E. coli* mutant, acting as "pathogen-like-bacteria," induces the UPR^mt^ as an intracellular surveillance mechanism for protection. We then established a forward genetics screen system using UPR^mt^ reporter P*hsp-6::gfp* which was activated by Δ*ymcB*, and found that neuronal MDSS-1 in ASE is required for sensing “pathogen-like-bacteria” and activating intestinal UPR^mt^ for against infection. MDSS-1,an uncharacterized single transmembrane protein expressed in ASE neuron, and the constitutive activation of MDSS-1 by overexpression with either the native promoter (Figure 5A) or a specific ASE neuronal promoter (Figure 5B) leads to the activation of intestinal UPR^mt^ in animals without Δ*ymcB* training, suggested that MDSS-1 acts as a potential receptor in neuron for sensing microbial signals and modulating mitochondria homoeostasis via cell-non-autonomous manner.

Cell-non-autonomous mitochondrial stress signal between neurons and intestine is a fitness strategy employed by worm to cope with various stress (Berendzen et al., 2016; Chen et al., 2021; Liu et al., 2022; Shao et al., 2016). Previous studies have identified several molecular mediators, such as neuronal peptides, Wnt/EGL-20, and GPCR, involved in the regulation of UPR^mt^ in a cell-non-autonomous manner. These studies utilized genetics methods, such as knocking out of mitochondrial genes (*cco-1, spg-7, atp-2*) in neuron, neuronal expression polyQ40, Wnt/EGL-20 and perturbing mitochondrial fusion in neurons by *fzo-1*mutation (Berendzen et al., 2016; Chen et al., 2021; Liu et al., 2022; Shao et al., 2016; Zhang et al., 2018) to study signal crosstalk between neurons and intestine. However, none of these methods identified the neuronal receptor MDSS-1 as a potential transducer of signals to the intestine for UPR^mt^ activation. There are two reasons, i) *mdss-1* expression is very low in ASE neuron, which cannot be detected via observing GFP level in transgenic knock-in animals by endogenous knocking in GFP to the MDSS-1 C terminus; 2) The intestinal UPR^mt^ activation by affecting neuronal mitochondrial damage is different of UPR^mt^ activation in animals through sensing bacteria. We provided evidence that the activation of intestinal UPR^mt^, induced by polyQ40 expression in neurons, remained unaffected by *mdss-1* mutations (Figure S3E). Therefore, our study also provides an efficiency research system to identify novel regulators in involved in regulation cell-non-autonomous mitochondrial stress. Understanding how mitochondrial stress is sensed in the nervous system and transmitted throughout the organism to induce UPR^mt^ in peripheral cells is crucial for the survival of *C. elegans* in nature, where precise sensing of bacteria is fundamental.

Collectively, this study identified specific *E. coli* mutants that behave like pathogenic bacteria and discovered a potential sensor in a specific neuron that modulates cell-non-autonomous mitochondrial surveillance in response to infection. This study also highlights the valuable system in identifying major mediators for mitochondrial surveillance in sensing bacteria by using native *C. elegans*-bacteria interaction system. In addition, our study also provides i) a roadmap for developing genetically engineered probiotics, ii) and potential way via targeting neuronal systems, in controlling intracellular surveillance or stress response programs for against infection or disease treatment.

### Limitations of the study

Our study reveals that MDSS-1 functions as a receptor in ASE neurons, enabling the detection of pathogen-like bacteria and triggering cell-non-autonomous UPR^mt^ as a defense mechanism against pathogens. Nevertheless, the specific molecular forms of “pathogen-like bacteria” that can be sensed by MDSS-1 remain unclear. Additionally, the mechanisms by which neuronal MDSS-1 influences the expression of endopeptidase inhibitors in the intestine are not yet fully understood, warranting further investigation.

## Author Contributions

H. L., P. C. and Y.X performed most experiments. F. H performed the biochemistry experiment. T.G performed some survival phenotype assay. Z.S guided some experiments and wrote/revised paper. B.Q. Z.S and H.L. wrote and edited the paper. B.Q. supervised this study.

## Acknowledgments

We thank the Caenorhabditis Genetics Center (CGC) (funded by NIH P40OD010440) for strains; Dr. Ye Tian for RNAi strains of *atfs-1* and stains of [P*rgef-1::ployQ40::YFP*;P*hsp-6::GFP*] and [P*rgef-1::egl-20*;P*hsp-6::GFP*]; Dr. Lingjun Zheng for RNAi strains of *mig-14;* This work was supported by the Ministry of Science and Technology of the People’s Republic of China (2019YFA0803100, 2019YFA0802100 to B. Q.), the National Natural Science Foundation of China (32071129 to Z.S; 32170794 to B.Q), Yunnan Provincial Science and Technology Project at Southwest United Graduate School (202302AP370005 to B.Q.), Yunnan Applied Basic Research Projects (202101AT070022, 202001AW070006 to Z.S., 202201AT070196 to B.Q), Yunnan Revitalization Talent Support Program (C619300A086 to Z.S., K264202230211 to B.Q.).

---

**Figure S1. Screen for "pathogen-like-bacteria" that protect animals against infection. Relative to Figure 1**.

**(A)** Cartoon illustration of *C. elegans* pre-treated with some natural bacterial species against pathogen infection.

**(B)** Cartoon illustration of a pathogen-like-bacteria pre-trained screening system in the laboratory. *C. elegans* pre-conditioned with some *E. coli* mutant strains enhance resistance against pathogen infection.

**(C)** Survival analysis of worms pre-conditioned with *E. coli K12* or mutant bacteria on PA14. Statistics are in Table S1.

For all panels, *p<0.05, **p<0.01, ***p<0.001, ****p<0.0001. All experiments were performed independently at least three times.

**Figure S2. Animals sense pathogen-like-bacteriaΔ*ymcB* through ATFS-1 depended UPR^mt^. Relative to Figure 2.**

**(A)** Quantification of selection for *E. coli* K12 or pathogen-like-bacteria of animals seeded into the middle of two bacteria for 60 hours at 20℃.

**(B)** Microscope images and fluorescence quantification of P*hsp-4::GFP,* P*hsp-6::GFP,*

P*irg-5::GFP* or P*sysm-1::GFP* reporter expression in wild-type animals grown on *E. coli K12 and* Δ*ymcB*.

**(C)** Microscope images and fluorescence quantification of P*hsp-6::GFP* reporter expression in wild-type animals feeding with either empty vector (EV), *atfs-1* (RNAi), *dve-1* (RNAi) or *ubl-5* (RNAi) followed by exposure to Δ*ymcB*.

For all panels, *p<0.05, **p<0.01, ***p<0.001, ****p<0.0001, n is the number of worms scored. Error bars, ± s.d. All experiments were performed independently at least three times.

**Figure S3. Transmembrane protein MDSS-1 is required for sensing pathogen-like-bacteria. Relative to Figure 3**.

**(A)** Cartoon illustration of established EMS screening system. EMS mutagenesis was performed in P*hsp-6::GFP* animals on Δ*ymcB* and mutant animals with effective mutations failed to be induced UPR^mt^, lacking enhanced pathogen resistance or preference phenotype.

**(B)** Microscope images and fluorescence quantification of P*hsp-6::GFP* reporter expression in animals feeding with empty vector (EV) or *mdss-1* (RNAi) followed by exposure to Δ*ymcB*.

**(C)** Microscope images and fluorescence quantification of P*hsp-6::GFP* reporter expression in *mdss-1(ylf14)* animals with or without P*rpl-28::mdss-1* transgene grown on Δ*ymcB*. Animals carrying transgenes discriminated by P*odr-1::GFP*.

**(D)** Fluorescence quantification of P*hsp-6::GFP* reporter expression in wild-type and *mdss-1(ylf14)* animals on pathogen-like-bacteria.

**(E)** Microscope images and fluorescence quantification of P*hsp-6::GFP* reporter expression in animals containing P*rgef-1::polyQ40::YFP* grown on empty vector (EV) or *mdss-1*(RNAi).

For all panels, *p<0.05, **p<0.01, ***p<0.001, ****p<0.0001, n is the number of worms scored. Error bars, ± s.d. All experiments were performed independently at least three times.

**Figure S4. The expression level of *mdss-1* under K12 or Δ*ymcB.* Relative to Figure 4**.

Microscope images and fluorescence quantification of P*mdss-1::mdss-1::mcherry* expression in animals grown on *E. coli* K12 and Δ*ymcB*. “n” is the number of worms scored. ns: no significant difference, n is the number of worms scored. Error bars, ± s.d. All experiments were performed independently at least three times.

**Figure S5. MDSS-1 acts as a potential receptor in UPR^mt^ signal transmission. Relative to Figure 5**.

**(A)** Microscope images and fluorescence quantification of *Phsp-6::GFP* reporter expression in animals containing P*mdss-1::mdss-1::mcherry* grown on either empty vector (EV), *atfs-1* (RNAi), *dve-1* (RNAi) or *ubl-5* (RNAi). Animals carrying transgenes discriminated by P*odr-1::GFP*.

**(B)** Microscope images and fluorescence quantification of P*hsp-6p::GFP* reporter expression in animals with or without P*gcy-6::mdss-1[Ex]::mcherry* transgene grown on *E. coli* K12.

**(C)** Microscope images and fluorescence quantification of *Phsp-6::GFP* in animals carrying P*vha-6::mdss-1::mcherry* transgene grown on either empty vector (EV), *atfs-1* (RNAi), *dve-1* (RNAi) or *ubl-5* (RNAi).

For all panels, *p<0.05, **p<0.01, ***p<0.001, ****p<0.0001, n is the number of worms scored. Error bars, ± s.d. All experiments were performed independently at least three times.

**Figure S6. The activation of MDSS-1-mediated UPR^mt^ involves the participation of inter-tissue communication factors. Relative to Figure 6**.

**(A)** Microscope images and fluorescence quantification of P*hsp-6p::GFP* reporter expression in animals containing P*mdss-1::mdss-1::mcherry* transgene grown on empty vector (EV), *flp-2* (RNAi) or *flp-5* (RNAi). Animals carrying transgenes discriminated by P*odr-1::GFP*.

**(B)** Microscope images and fluorescence quantification of P*hsp-6p::GFP* reporter expression in animals containing P*mdss-1::mdss-1::mcherry* grown on empty vector (EV), *egl-6* (RNAi) or *npr-42* (RNAi). Animals carrying transgenes discriminated by P*odr-1::GFP*.

**(C)** Microscope images of wild-type animals carrying P*nlp-3::nlp-3::mcherry* and P*gcy-6::GFP*.

**(D)** Microscope images and fluorescence quantification of *Phsp-6p::GFP* reporter expression in animals containing P*mdss-1::mdss-1::mcherry* transgene grown on either empty vector (EV), *fshr-1* (RNAi) or *srz-75* (RNAi). Animals carrying transgenes discriminated by P*odr-1::GFP*.

**(E)** Microscope images and fluorescence quantification of P*hsp-6p::GFP* reporter expression in animals containing P*mdss-1::mdss-1::mcherry* transgene grown on either empty vector (EV), *mig-14* (RNAi) or *egl-20* (RNAi). Animals carrying transgenes discriminated by P*odr-1::GFP*.

**(F)** Microscope images and fluorescence quantification of P*hsp-6::GFP* reporter expression in animals containing *Prgef-1::egl-20* transgene grown on empty vector (EV) and *mdss-1*(RNAi).

For all panels, *p<0.05, **p<0.01, ***p<0.001, ****p<0.0001, n is the number of worms scored. Error bars, ± s.d. All experiments were performed independently at least three times.

**Figure S7. The activation of MDSS-1-mediated UPR^mt^ involves the endopeptidase inhibitors. Relative to Figure 6**.

**(A)** Heatmap of deferentially down-regulated genes induced by Δ*ymcB* in group I

(wild-type animals grown on K12 followed by exposure to K12 or Δ*ymcB*) and group II (*mdss-1(ylf14)* animals grown on K12 followed by exposure to K12 or Δ*ymcB*). Genes with an adjusted p value< 0.05 were selected as differentially expressed genes.

**(B)** Venn diagram of numbers of deferentially down-regulated genes in wild-type and *mdss-1(ylf14)* mutant animals grown on K12 transferred to Δ*ymcB* for 24 hours. Genes with an adjusted p value< 0.05 were selected as deferentially expressed genes.

**(C)** GO enrichment of overlapped genes in Figure 7(D). Gene numbers are shown in circle size and P.values are shown in color. See also table S4.

**(D)** A schematic of CLPP-1 involved UPR^mt^ in *C.elegans*. The activity of CLPP-1 that degrading the unfolded protein into peptide fragments was inhibited by serine-type endopeptidase inhibitors, which in turn affected the activation of UPR^mt^ downstream.

**(E)** The genes with serine-type endopeptidase inhibitor activity, which are negatively regulated by MDSS-1, were identified through RNA-seq analysis from Group II.

**(F)** Heatmap Score of these endopeptidase inhibitors in different tissues. Low score is shown in white, and high score is in blue. See also Table S5. Data from worm tissue prediction application: https://worm.princeton.edu/ (Kaletsky R et al., 2018)

**Table S1.** Statistics for survival analysis. Related to Figure 1B, 2F, 3F, 5C, 5D and Figure S1C.

**Table S2.** List of screened neuropeptides. Related to Figure 6 and Figure S6.

**Table S3.** Deferentially genes expressed in group I (wild-type animals grown on K-12 transferred to K-12 and Δ*ymcB* respectively) and group II (*mdss-1(ylf14)* animals grown on K12 transferred to K12 and Δ*ymcB* respectively). Related to Figure 6D-E and Figure S7A-B

**Table S4.** Expression and GO enrichment of overlapped genes in Figure 6E and S7C. Related to Figure 6E and Figure S7C.

**Table S5.** Heatmap score of inhibitor expression in different tissue. Related to Figure S7F. Data download from https://worm.princeton.edu/ (Kaletsky R et al., 2018).

**Table S6.** List of Oligonucleotide Sequences, Related to STAR Methods Key Resource Table.

## Star Methods

### KEY RESOURCES TABLE

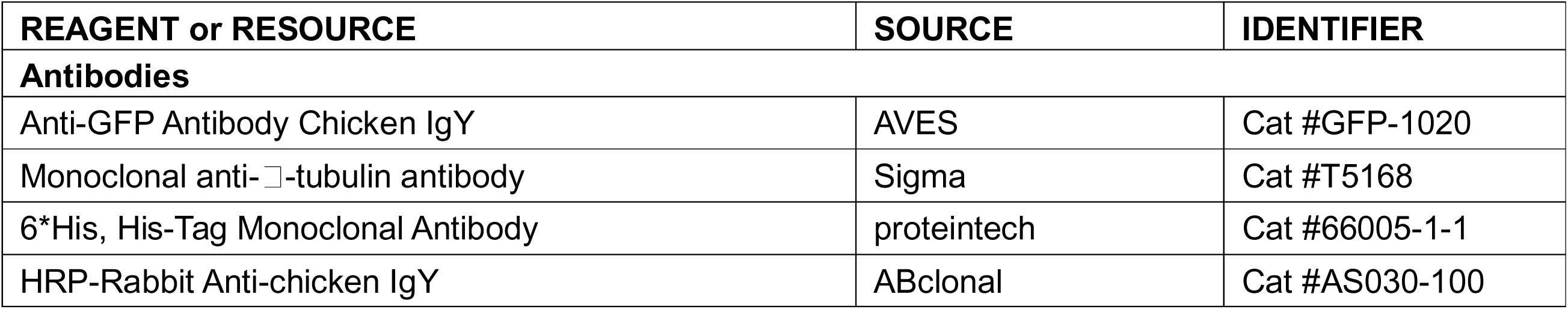
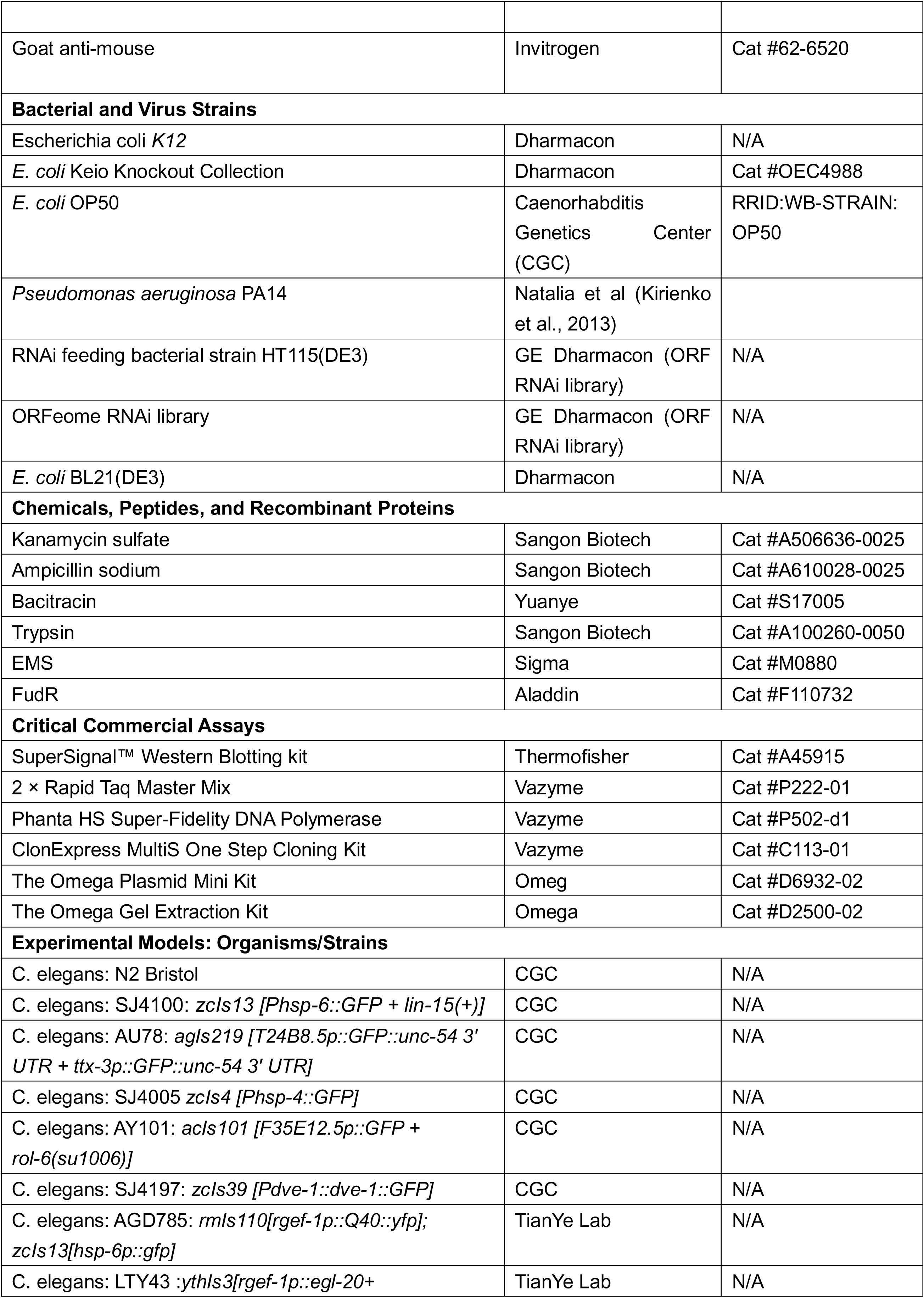
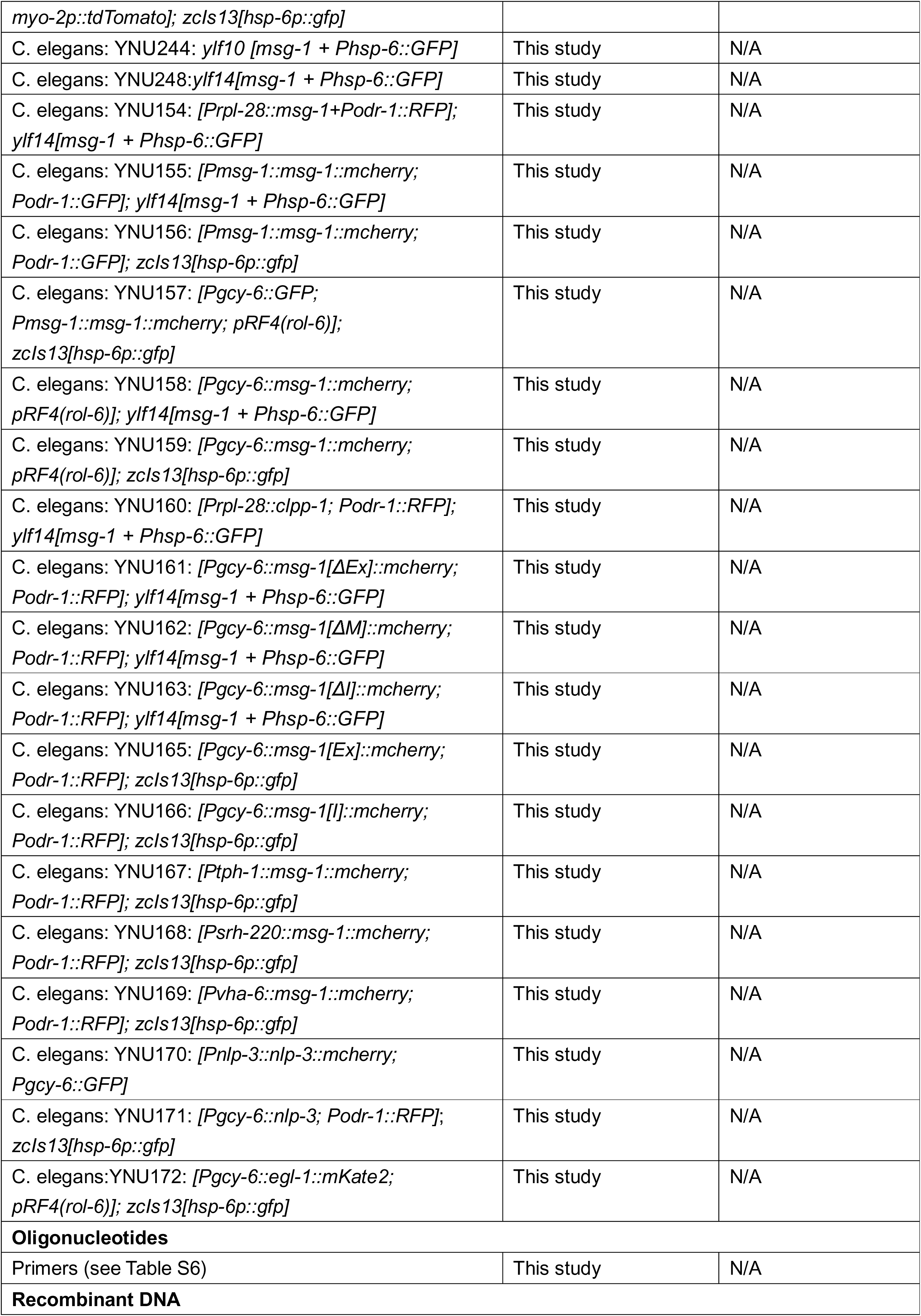
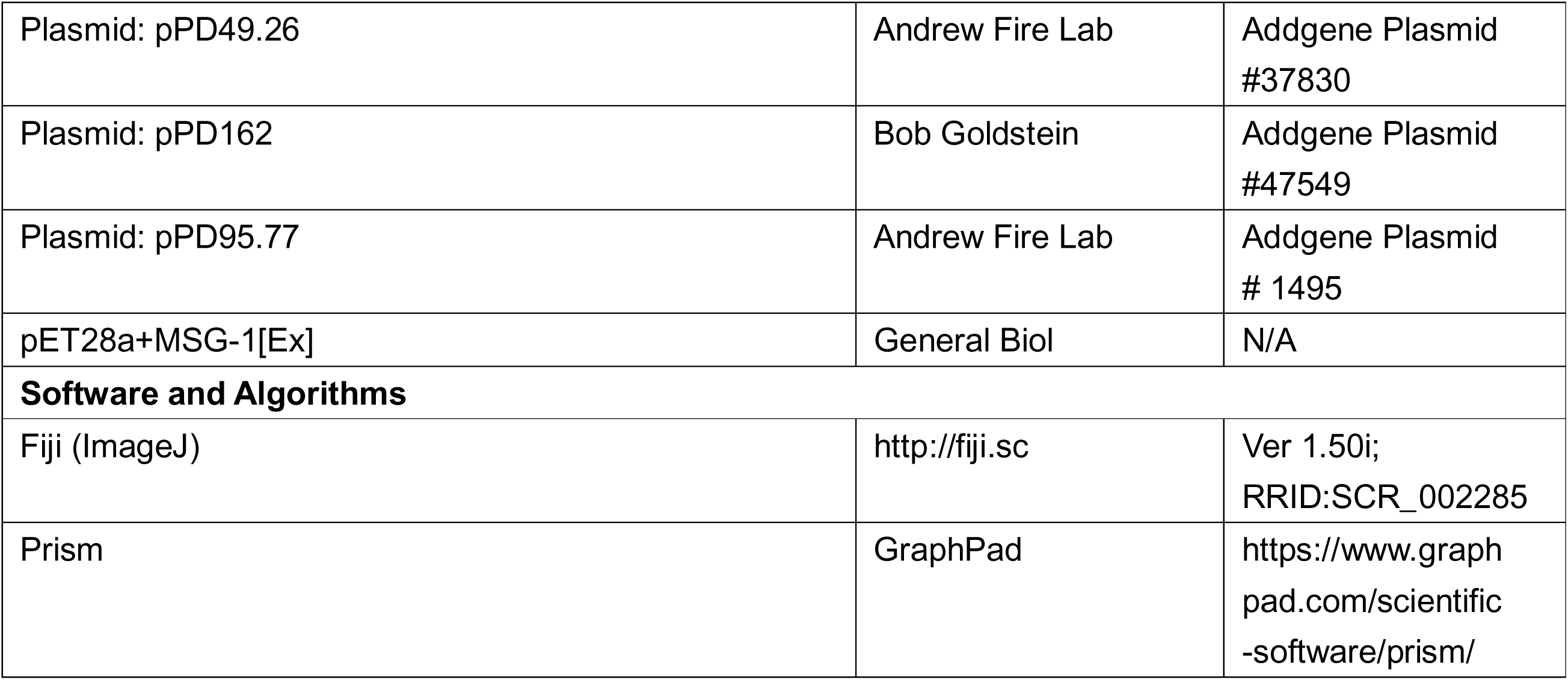

### Resource Availability

#### Lead contact

Further information and requests for reagents may be directed to the Lead contact Bin Qi.

#### Materials availability

All reagents and strains generated by this study are available through request to the lead contact with a completed Material Transfer Agreement.

#### Data and code availability

RNA-seq data are accessible in Table S3. This paper does not report original code.

Any additional information required to reanalyze the data reported in this paper is available from the lead contact upon request.

### Experimental model and subject details

#### *C. elegans* strains and maintenance

Nematode stocks were maintained on nematode growth medium (NGM) plates seeded with bacteria (*E. coli* OP50) at 20 °C.

1) The following strains were obtained from the Caenorhabditis Genetics Center (CGC):

N2 Bristol (wild-type); SJ4100: *zcIs13 [Phsp-6::GFP + lin-15(+)]*; AU78: *agIs219*

*[T24B8.5p::GFP::unc-54 3’ UTR + ttx-3p::GFP::unc-54 3’ UTR]*; SJ4005: *zcIs4 [Phsp-4::GFP]*; AY101: *acIs101 [F35E12.5p::GFP + rol-6(su1006)]*; SJ4197: *zcIs39 [Pdve-1::dve-1::GFP]*.

2) The following strains were obtained from TianYe Lab:

AGD785: *rmIs110*[P*rgef-1::Q40::yfp*]; *zcIs13*[*hsp-6p::gfp*] LTY43:*ythIs3*[P*rgef-1::egl-20*+ *myo-2p::tdTomato*]; *zcIs13*[*hsp-6p::gfp*]

3) The following strains were constructed in this study:

YNU244: *ylf10 [mdss-1 +* P*hsp-6::GFP]* was obtained from EMS mutagenesis in SJ4100 background.

YNU248: *mdss-1(ylf14)* terminating mutant was constructed by Crispr/Cas9 in SJ4100 background.

YNU154: [P*rpl-28::mdss-1;* P*odr-1::RFP*] transgene strain was constructed by injecting plasmid P*rpl-28::mdss-1* with P*odr-1::RFP* in YNU248 background.

YNU155: [P*mdss-1::mdss-1::mcherry*; P*odr-1::GFP*] transgene strain was constructed by injecting plasmid P*mdss-1::mdss-1::mcherry* with P*odr-1::GFP* in YNU248 background. YNU156: [P*mdss-1::mdss-1::mcherry*; P*odr-1::GFP*] transgene strain was constructed by injecting plasmid P*mdss-1::mdss-1::mcherry* with P*odr-1::GFP* in SJ4100 background.

YNU157: [P*gcy-6::GFP*; P*mdss-1::mdss-1::mcherry*; pRF4*(rol-6)*] transgene strain was constructed by injecting plasmid P*gcy-6::RFP*, P*mdss-1::mdss-1::mcherry* and pRF4*(rol-6)*in SJ4100 background.

YNU158: [P*gcy-6::mdss-1::mcherry*; pRF4*(rol-6)*] transgene strain was constructed by injecting plasmid P*gcy-6::mdss-1::mcherry* with pRF4*(rol-6)* in YNU248 background. YNU159: [P*gcy-6::mdss-1::mcherry*; pRF4*(rol-6)*] transgene strain was constructed by injecting plasmid P*gcy-6::mdss-1::mcherry* with pRF4*(rol-6)* in SJ4100 background.

YNU160: [P*rpl-28::clpp-1*; P*odr-1::RFP*] transgene strain was constructed by injecting plasmid P*rpl-28::clpp-1* with P*odr-1::RFP* in YNU248 background.

YNU161: [P*gcy-6::mdss-1[*Δ*Ex]::mcherry*; P*odr-1::RFP*] transgene strain was constructed by injecting plasmid P*gcy-6::mdss-1[*Δ*Ex]::mcherry* with P*odr-1::RFP* in YNU248 background.

YNU162: [P*gcy-6::mdss-1[*Δ*M]::mcherry*; P*odr-1::RFP*] transgene strain was constructed by injecting plasmid P*gcy-6::mdss-1[*Δ*M]::mcherry* with P*odr-1::RFP* in YNU248 background.

YNU163: [P*gcy-6::mdss-1[*Δ*I]::mcherry*; P*odr-1::RFP*] transgene strain was constructed by injecting plasmid P*gcy-6::mdss-1[*Δ*I]::mcherry* with P*odr-1::RFP* in YNU248 background.

YNU165: [P*gcy-6::mdss-1[Ex]::mcherry*; P*odr-1::RFP*] transgene strain was constructed by injecting plasmid P*gcy-6::mdss-1[Ex]::mcherry* with P*odr-1::RFP* in SJ4100 background.

YNU166: [P*gcy-6::mdss-1[I]::mcherry*; P*odr-1::RFP*] transgene strain was constructed by injecting plasmid P*gcy-6::mdss-1[I]::mcherry* with P*odr-1::RFP* in SJ4100 background.

YNU167: [P*tph-1::mdss-1::mcherry*; P*odr-1::RFP*] transgene strain was constructed by injecting plasmid P*tph-1::mdss-1::mcherry* with P*odr-1::RFP* in SJ4100 background.

YNU168: [P*srh-220::mdss-1::mcherry*; P*odr-1::RFP*] transgene strain was constructed by injecting plasmid P*srh-220::mdss-1::mcherry* with P*odr-1::RFP* in SJ4100 background.

YNU169: [P*vha-6::mdss-1::mcherry*; P*odr-1::RFP*] transgene strain was constructed by injecting plasmid P*vha-6::mdss-1::mdss-1::mcherry* with P*odr-1::RFP* in SJ4100 background.

YNU170: [P*nlp-3::nlp-3::mcherry*;P*gcy-6::GFP*] transgene strain was constructed by injecting plasmids P*nlp-3::nlp-3::mcherry* and P*gcy-6::GFP* in N2 background.

YNU171: [P*gcy-6::nlp-3*;P*odr-1::RFP*] transgene strain was constructed by injecting plasmid P*gcy-6::nlp-3::GFP* with P*odr-1::RFP* in SJ4100 background.

YNU172: [P*gcy-6::egl-1::mKate2*;pRF4(*rol-6*)] transgene strain was constructed by injecting plasmid P*gcy-6::egl-1::mKate2* with pRF4(*rol-6*) in SJ4100 background.

### Bacterial strains

*E. coli-*OP50, *E. coli-*K12 (BW25113), *E. coli-*K12 mutant, *P.aeruginosa* were cultured at 37°C in LB medium. A standard cultured bacteria (OD600=1) was then spread onto each Nematode growth media (NGM) plate.

## Method Details

### Generation of transgenes

1. To construct the *C. elegans* plasmid for expression of mdss-1 in multiple tissues, 1437bp promoter of *rpl-28* and genomic DNA of *mdss-1* was cloned into the pPD95.77 vector. DNA plasmid mixture containing P*rpl-28::mdss-1::GFP*(20ng/μL) and P*odr-1p:RFP*(50ng/μL) was injected into the gonads of adult *mdss-1(ylf14)* animals.
2. To construct the *C. elegans* plasmid for expression of *mdss-1* in its native tissue, 495bp promoter and genomic DNA of *mdss-1* was cloned into the pPD95.77 vector. DNA plasmid mixture containing P*mdss-1::mdss-1::RFP* (20ng/μL) and P*odr-1p::GFP*(50ng/μL) was injected into the gonads of adult *mdss-1(ylf14)* and wild-type animals.
3. To construct the *C. elegans* plasmid for expression of *mdss-1* in the ASE neurons, 2000bp promoter of *gcy-6* and genomic DNA of *mdss-1* was cloned into the pPD95.77 vector. DNA plasmid mixture containing P*gcy-6::mdss-1::mcherry* (20ng/μL) and pRF4*(rol-6)* (50ng/μL) was injected into the gonads of adult *mdss-1(ylf14)* and wild-type animals.
4. To construct the *C. elegans* plasmid for expression of *mdss-1* isoforms with varying lengths in ASE neuron, 561bp genomic DNA of *mdss-1[*Δ*Ex]*, 1723bp genomic DNA of *mdss-1[*Δ*M]*, and 1788bp genomic DNA of *mdss-1[*Δ*I]* in conjunction with 2000bp promoter of *gcy-6* was cloned into the pPD95.77 vector. DNA plasmid mixture containing either P*gcy-6::mdss-1[*Δ*Ex]::mcherry*(20ng/μL), P*gcy-6::mdss-1[*Δ*M]::mcherry*(20ng/μL) or P*gcy-6::mdss-1[*Δ*I]::mcherry*(20ng/μL) and P*odr-1::RFP*(50ng/μL) injected into the gonads of adult *mdss-1(ylf14)* animals.
5. To construct the *C. elegans* plasmid for overexpression of *mdss-1* isoforms with varying lengths in ASE neuron, 1225bp genomic DNA of *mdss-1[Ex]* and 498bp genomic DNA of *mdss-1[I]* in conjunction with 2000bp promoter of *gcy-6* was cloned into the pPD95.77 vector. DNA plasmid mixture containing either P*gcy-6::mdss-1[Ex]::mcherry*(20ng/μL) or P*gcy-6::mdss-1[I]::mcherry*(20ng/μL) and P*odr-1::RFP*(50ng/μL) injected into the gonads of adult wild-type animals.
6. To construct the *C. elegans* plasmid for expression of *mdss-1* in either ADF neuron, ADL neuron or intestine, 3108bp promoter of *tph-1*, 4003bp promoter of *srh-220* or 1593bp promoter of *vha-6* and genomic DNA of *mdss-1* was cloned into the pPD95.77 vector. DNA plasmid mixture containing either P*tph-1::mdss-1::mcherry*(20ng/μL), P*srh-220::mdss-1::mcherry*(20ng/μL) or P*vha-6::mdss-1::mcherry*(20ng/μL) and P*odr-1::RFP*(50ng/μL) injected into the gonads of adult wild-type animals.
7. To construct the *C. elegans* plasmid for expression of *clpp-1*, 1437bp promoter of *rpl-28* and genomic DNA of *clpp-1* was cloned into the pPD95.77 vector. DNA plasmid mixture containing P*rpl-28::clpp-1*(20ng/μL) and P*odr-1p:RFP*(50ng/μL) was injected into the gonads of adult *mdss-1(ylf14)* animals.
8. To construct the *C. elegans* plasmid for co-colonization of expression of *gcy-6* and *mdss-1*, 2000bp promoter of *gcy-6* was cloned into the pPD95.77 vector. DNA plasmid mixture containing P*gcy-6::GFP* (20ng/μL), P*mdss-1::mdss-1::mcherry* (20ng/μL) and pRF4(rol-6) (50ng/μL) was injected into the gonads of SJ4100 animals.
9. To construct the *C. elegans* plasmid for co-colonization of expression of *gcy-6* and *nlp-3*, 2000bp promoter of nlp-3 and genomic DNA was cloned into the pPD49.26 vector, 2000 promoter of gcy-6 was cloned into the pPD95.77 vector. DNA plasmid mixture containing P*nlp-3::nlp-3::mcherry*(20ng/μL) and P*gcy-6p:GFP*(50ng/μL) was injected into the gonads of adult wild-type animals.
10. To construct the *C. elegans* plasmid for expression of *nlp-3* in ASE neuron, 2000bp promoter of *gcy-6* and genomic DNA of nlp-3 was cloned into the pPD95.77 vector. DNA plasmid mixture containing P*gcy-6::nlp-3*(20ng/μL) and P*odr-1p:RFP*(50ng/μL) was injected into the gonads of adult wild-type animals.
11. To construct the *C. elegans* plasmid for the ablation of ASE neuron, 2000bp promoter of *gcy-6* and genomic DNA of egl-1 was cloned into the pPD49.26 vector. DNA plasmid mixture containing P*gcy-6::egl-1::mKate2*(20ng/μL) and pRF4(*rol-6*)(50ng/μL) was injected into the gonads of adult wild-type animals.

### CRISPR/Cas9 mediated gene editing

To generate *mdss-1(ylf14)* stop gained mutants, the two sgRNA targeting sequence (5’-catcgtggcagtgagaatg-3’,5’-cattctcactgccacgatg-3’) designed in close proximity to the start codon of *mdss-1* were cloned into the pDD162 vector. A repair template containing the enzymatic cut site of the terminating mutation was injected along with the sgRNA vectors and a co-marker plasmid (P*odr-1::RFP*) into wild-type animals. The micro-injection system was composed by 2μM repair template,50ng/μL co-marker, and 20ng/μL sgRNA vectors. Heterozygous F1 animals were obtained based on co-marker expression, and the subsequent F2 generation was screened via PCR to identify mutant animals.

### “Pathogen-like-bacteria” screening

Overnight incubation of bacteria of *E. coli* K12 Keio library in LB liquid in 96-well plates and the addition of 50μg/mL kanamycin is necessary. *E. coli* mutants are from the Keio *E. coli* single mutant collection (Baba et al., 2006). Mutant bacteria strains, as well as the wild-type control strain BW25113, were cultured overnight in LB medium with 50 ug/ml kanamycin in 96-well plates at 37°C. The cultured bacteria was spread onto 35mm NGM plates. The synchronized L1 worms were then seeded onto plates and grown at 20 °C until to adulthood. Meanwhile PA14 was cultured at 37 °C overnight and then the concentrated tenfold culture was spread over 35mm NGM medium with 50mg/L FudR. PA14 plates were cultured at 37 °C 24 hours then placed at 25 °C 24 hours for preparation. Adult animals on K12 or mutant bacteria were transferred to PA14 plates (50 worms per plate). In this study, we tested 4320 different mutant strains of *E. coli* to see if any of them could protect animals from PA14 infection. Only the mutants that showed higher survival rates than the control group at a specific time point were chosen for further analysis. As a result, we identified 20 *E. coli* mutants that provided protection against PA14 infection. We have provided detailed information about how these mutants performed in terms of animal survival when exposed to PA14 in Figure 1B, Figure S1B and Table S1.

### "Pathogen-like-bacteria" selection assays

The food choice assay was modified from Qi et al (Qi et al., 2017). Briefly, 10µL of K12 and "pathogen-like-bacteria" was added to indicated position of 35mm NGM plate (Figure S2A). Synchronized L1 nematodes were seeded onto the center. The number of animals were counted after 60 hours on 20℃.

### *P. aeruginosa* killing assays

The method was modified from Tan et al (Tan et al., 1999). Briefly, 100μL of overnight culture PA14 was spread over 35mm NGM plates with 50mg/L FUdR. PA14 plates were cultured at 37 °C 24 hours then placed at 25 °C 24 hours for preparation. Adult animals were transferred to PA14 plates. 50 animals were transferred per plate, and three replicate plates were necessary. The number of deaths was counted every 12 hours and the carcasses were picked out. After the count was completed, the number of deaths was added up as the denominator for the survival rate calculation.

### EMS mutagenesis

Synchronized L1 animals were grown until the L4 stage on Δ*ymcB* and were then exposed to a 4-hour treatment of 0.5% EMS (Ethyl Methanesulfonate). Following treatment, the animals were thoroughly washed with M9 solution and transferred to Δ*ymcB* plates. The P0 generation, consisting of approximately 1000 worms, was distributed onto multiple Δ*ymcB* plates to ensure continuous reproduction without starvation. Every 3-4 F1 animals were selectively picked from a Δ*ymcB* plate (3-4 F1 worms per plate) to generate F2 through self-fertilization. A total of 3000 plates containing F1 animals (approximately 9000-12000 F1 worms) were screened in order to identify F2 animals with low P*hsp-6::GFP* expression. To confirm the suspected mutant, it was picked singly onto Δ*ymcB* for passaging. During the screen, we discovered four mutant animals that exhibited significantly reduced P*hsp-6::GFP* expression under Δ*ymcB* feeding conditions. However, we were especially intrigued by one particular mutant called *mdss-1*, as it is a gene that is only expressed in the neuron.

### Identification of EMS mutants

DNA isolation, library construction, and whole genome sequencing with gene identified were carried out according to the published protocol (Joseph et al., 2018).

#### **i)** Preparation of samples for Whole-Genome-Sequencing (WGS)

For each backcross, wild-type males were crossed to mutant hermaphrodites on Δ*ymcB* feeding condition, F1 cross-progeny animals were individually cloned and F2 variants and wild-types were isolated. The variant strains and wild-type strains were mixed respectively to constitute the “DNA-pool” used as samples for WGS.

Whole-genome sequencing of pooled F2 recombinants, homozygous for the mutant phenotype following two outcrosses to wild-type N2 animals, was performed to identify the mutations that caused constitutive *Phsp-6::GFP* expression.

#### **ii)** WGS data processing

For Whole-Genome-Sequencing, paired-end libraries were sequenced on an Illumina HiSeq 2000. Fastqc was used to control the quality of raw data and Trimmomatic was used to filter the data. Bwa was used to construct the index of *C.elegans* genome and align clean reads to the reference gene sequence (Species: *Caenorhabditis_elegans*; Source: UCSC; Reference Genome Version: WBcel235/ce11). The Samtools was used for file format conversion and sorting. Picard was used to remove the duplicate reads and then GATK was used to identify the intervals and realign. The realigned sequence was piled up by Samtools and then inputted to Varscan to call the variants including SNPs and INDELs. Vcflib was used to perform subtraction between the wild-types and variants. The file was annotated by Snpeff and the candidate genes was finally obtained.

### RNAi feeding

All RNAi by feeding used bacterial clones from the MRC RNAi library (Kamath et al., 2003) or the ORF-RNAi Library (Rual et al., 2004). RNAi plates were prepared by adding 100μg/mL ampicillin and 1mM IPTG to NGM agar. And overnight cultures of specific RNAi strains and the control HT115 strain with 100μg/mL ampicillin were seeded onto RNAi feeding plates. Synchronized L1 wild-type animals grown on the RNAi plates to the L4-stage were transferred to the Δ*ymcB* or K12 plates with 50μg/mL kanamycin, phenotypic observation occurred 24 hours after transfer.

### Preparation of samples for RNA-Sequencing

RNA-seq was done with three biological replicates that were independently generated, collected, and processed.

1. For the wild-type animals samples preparation, synchronized L1 wild-type animals seeded onto K12 plates grown to L4-stage were transferred to K-12 and Δ*ymcB* plates respectively, the samples were collected after 24 hours.
2. For the *mdss-1(ylf14)* mutant samples preparation, synchronized L1 *mdss-1(ylf14)* animals seeded onto K12 plates grown to L4-stage were transferred to K12 and Δ*ymcB* plates respectively plates, the samples were collected after 24 hours.

### Western blot

To quantify the protein of expression Hsp-6p*::GFP*, wild-type and mutant animals treated differently were analyzed by standard western blot methods and probed with anti-GFP (dilution = 1:5,000; AVES, GFP-1020) and anti-chicken IgY (dilution = 1:10,000; ABclonal, AS030) or anti-tubulin (dilution =1:10,000; Sigma T5168) as a loading control.

### Measurement of the fluorescence intensity

For fluorescence images of the UPR^mt^ phenotype, nematodes were anesthetized with 10mM levamisole and photographed under excitation light. The fluorescence intensity of whole worm was counted by ImageJ except for the transgenic nematodes expressing RFP/GFP co-marker in the nerve and its control group where only intestinal fluorescence was counted. The images of DVE-1::GFP aggregation taken at maximum magnification in a stereomicroscope and quantified by numbers of puncta.

## QUANTIFICATION AND STATISTICAL ANALYSIS

### Quantification

Transgenic animals were randomly selected for fluorescent photography. And then ImageJ software was used for quantifying fluorescence intensity of P*hsp-6::gfp* reporter which was normalized to body area.

### Statistical analysis

Except for survival curves in the *P. aeruginosa* killing assays were plotted by Prism and the significant difference of two groups of samples was analyzed by Log-rank (Mantel-Cox) test, two-tailed unpaired t-test was used for statistical analysis of two columns of samples in the rest of assays. Using the mean value of the control group as the baseline, the fold change in multiple of all individual values relative to the baseline was taken as the “Relative Intensity”. The error line in the data visualization is indicated by Mean±SD and the p-value representing significant difference is shown by “*”. For all figures, the numbers of animals scored were shown by “n”. And Venny 2.1.0 was used to analyze the overlap of gene sets. All experiments were performed independently at least three times, except for the original mutated *E. coli* screen.

