## Supplemental Figures for "Bacterial sensing via Neuronal Receptor Initiates Gut Mitochondrial Surveillance for Host Adaptation"

### Figure S1

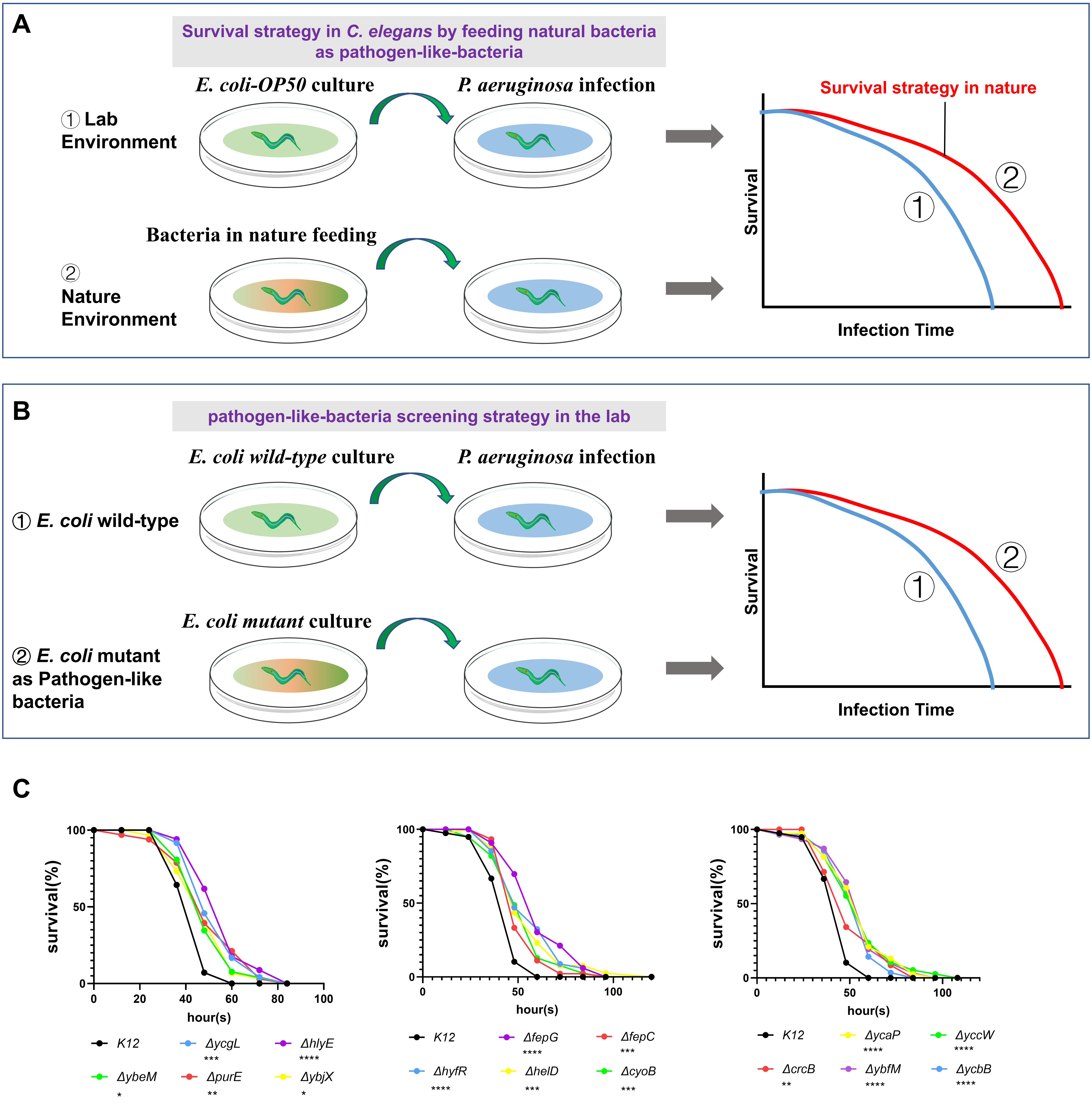


**Figure S1. Establishment of "pathogen-like-bacteria" screening strategy in *C. elegans* training with *E. coli* mutant. Relative to Figure 1.**
(A) Cartoon illustration of *C. elegans* pre-treated with some natural bacterial species against pathogen infection.
(B) Cartoon illustration of a pathogen-like-bacteria pre-trained screening system in the laboratory. *C. elegans* pre-conditioned with some *E. coli* mutant strains enhance resistance against pathogen infection.
(C) Survival analysis of worms pre-conditioned with *E. coli K12* or mutant bacteria on PA14. Statistics are in Table S1. . Statistics are in Table S1.
For all panels, *p<0.05, **p<0.01, ***p<0.001, ****p<0.0001. All experiments were performed independently at least three times.

### Figure S2

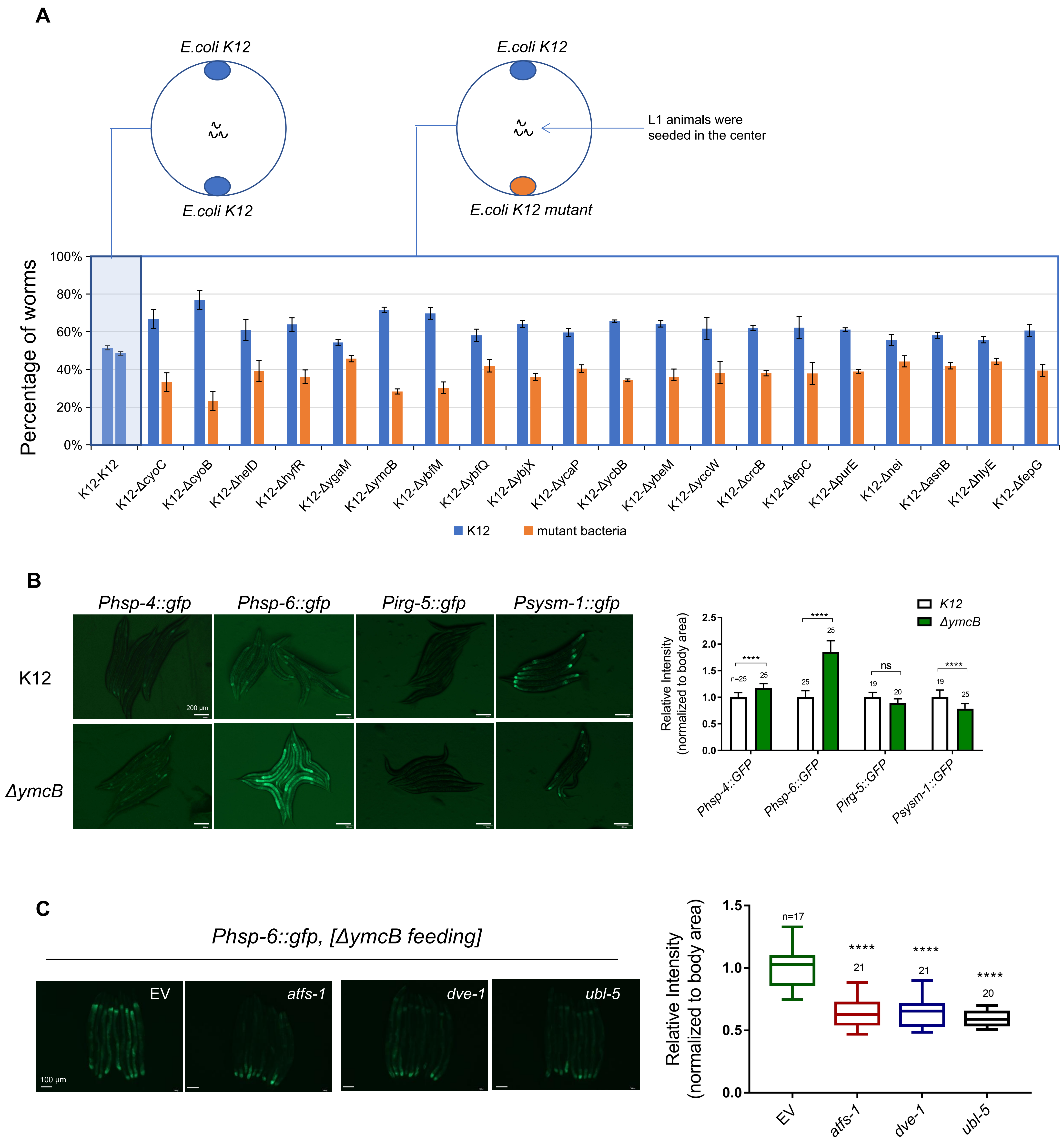


**Figure S2. Animals sense pathogen-like-bacteria *ΔymcB* through ATFS-1 depended UPR$^{mt}$. Relative to Figure 2.**
(A) Quantification of selection for *E. coli* K12 or pathogen-like-bacteria of animals seeded into the middle of two bacteria for 60 hours at 20℃.
(B) Microscope images and fluorescence quantification of P*hsp-4::GFP,* P*hsp-6::GFP,* P*irg-5::GFP* or P*sysm-1::GFP* reporter expression in wild-type animals grown on *E. coli K12 and ΔymcB*.
(C) Microscope images and fluorescence quantification of P*hsp-6::GFP* reporter expression in wild-type animals feeding with either empty vector (EV), *atfs-1* (RNAi), *dve-1* (RNAi) or *ubl-5* (RNAi) followed by exposure to *ΔymcB*.
For all panels, *p<0.05, **p<0.01, ***p<0.001, ****p<0.0001, n is the number of worms scored. Error bars, ± s.d. All experiments were performed independently at least three times.

### Figure S3

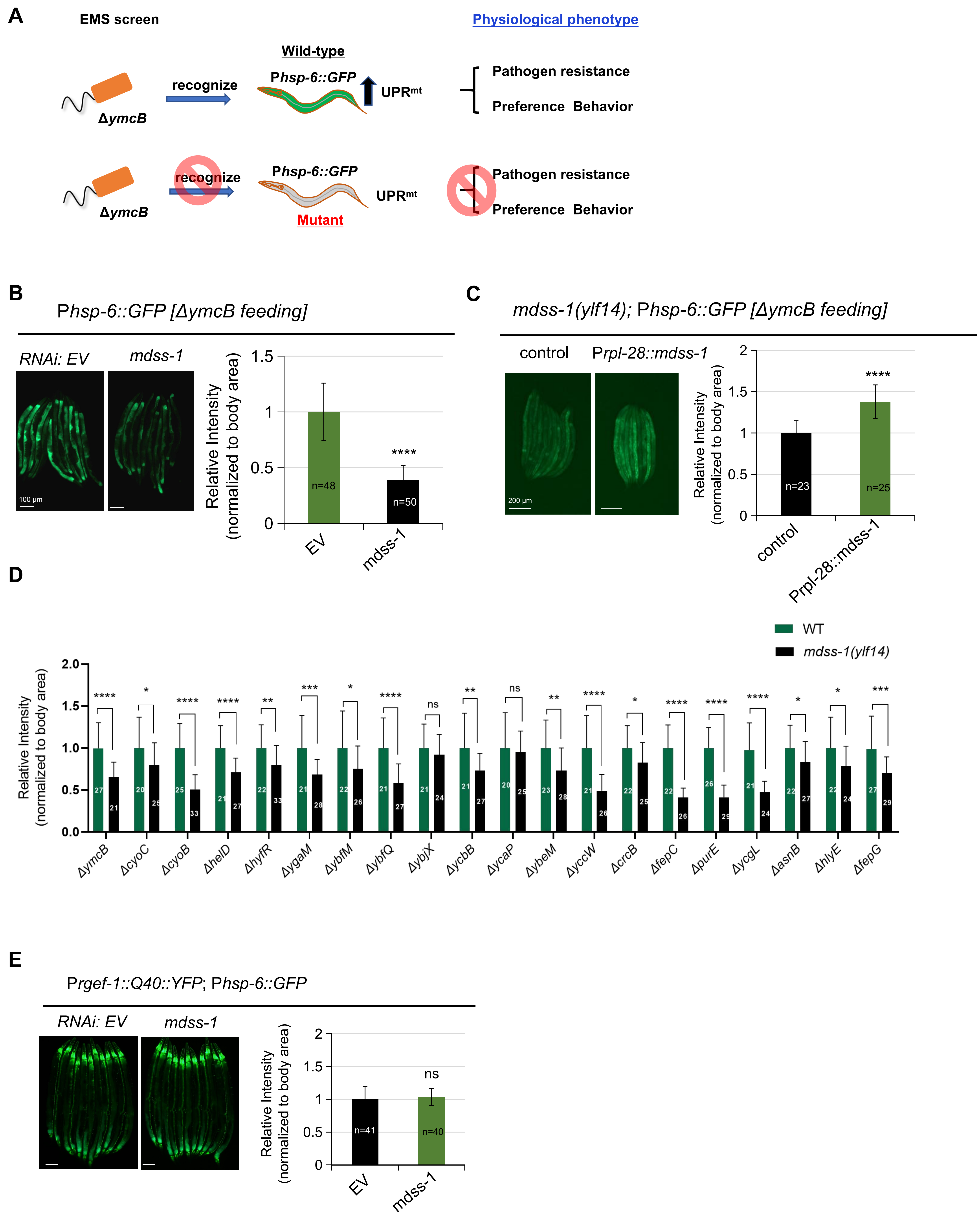

A
EMS screen
Physiological phenotype
Wild-type
ΔymcB
recognize
Phsp-6::GFP
UPRmt
Pathogen resistance
Preference Behavior
Mutant
B
Phsp-6::GFP [ΔymcB feeding]
RNAi: EV
mdss-1
100 μm
Relative Intensity (normalized to body area)
n=48
n=50
****
EV
mdss-1
C
mdss-1(ylf14); Phsp-6::GFP [ΔymcB feeding]
control
Prpl-28::mdss-1
200 μm
n=23
n=25
D
WT
mdss-1(ylf14)
ΔymcB
ΔcyoC
ΔcyoB
ΔhelD
ΔhyfR
ΔygaM
ΔybfM
ΔybfQ
ΔybjX
ΔycbB
ΔycaP
ΔybeM
ΔyccW
ΔcrcB
ΔfepC
ΔpurE
ΔycgL
ΔasnB
ΔhlyE
ΔfepG
E
Prgef-1::Q40::YFP; Phsp-6::GFP
n=41
n=40
ns

**Figure S3. Transmembrane protein MDSS-1 is required for sensing pathogen-like-bacteria. Relative to Figure 3.**

**(A)** Cartoon illustration of established EMS screening system. EMS mutagenesis was performed in P*hsp-6::GFP* animals on *ΔymcB* and mutant animals with effective mutations failed to be induced $UPR^{mt}$, lacking enhanced pathogen resistance or preference phenotype.

**(B)** Microscope images and fluorescence quantification of P*hsp-6::GFP* reporter expression in animals feeding with empty vector (EV) or *mdss-1* (RNAi) followed by exposure to *ΔymcB*.

**(C)** Microscope images and fluorescence quantification of P*hsp-6::GFP* reporter expression in *mdss-1(ylf14)* animals with or without P*rpl-28::mdss-1* transgene grown on *ΔymcB*. Animals carrying transgenes discriminated by P*odr-1::GFP*.

**(D)** Fluorescence quantification of P*hsp-6::GFP* reporter expression in wild-type and *mdss-1(ylf14)* animals on pathogen-like-bacteria.

**(E)** Microscope images and fluorescence quantification of P*hsp-6::GFP* reporter expression in animals containing P*rgef-1::polyQ40::YFP* grown on empty vector (EV) or *mdss-1*(RNAi).

For all panels, *p<0.05, **p<0.01, ***p<0.001, ****p<0.0001, n is the number of worms scored. Error bars, ± s.d. All experiments were performed independently at least three times.

### Figure S4

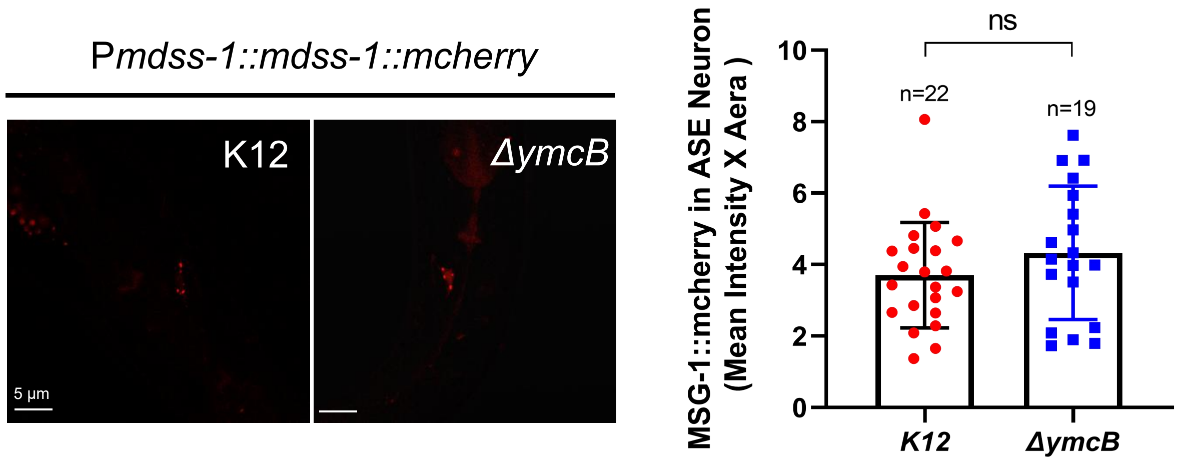


**Figure S4. The expression level of *mdss-1* under K12 or *ΔymcB.* Relative to Figure 4.**
Microscope images and fluorescence quantification of P*mdss-1::mdss-1::mcherry* expression in animals grown on *E. coli* K12 and *ΔymcB*. ns: no significant difference , n is the number of worms scored. Error bars, ± s.d. All experiments were performed independently at least three times.

### Figure S5

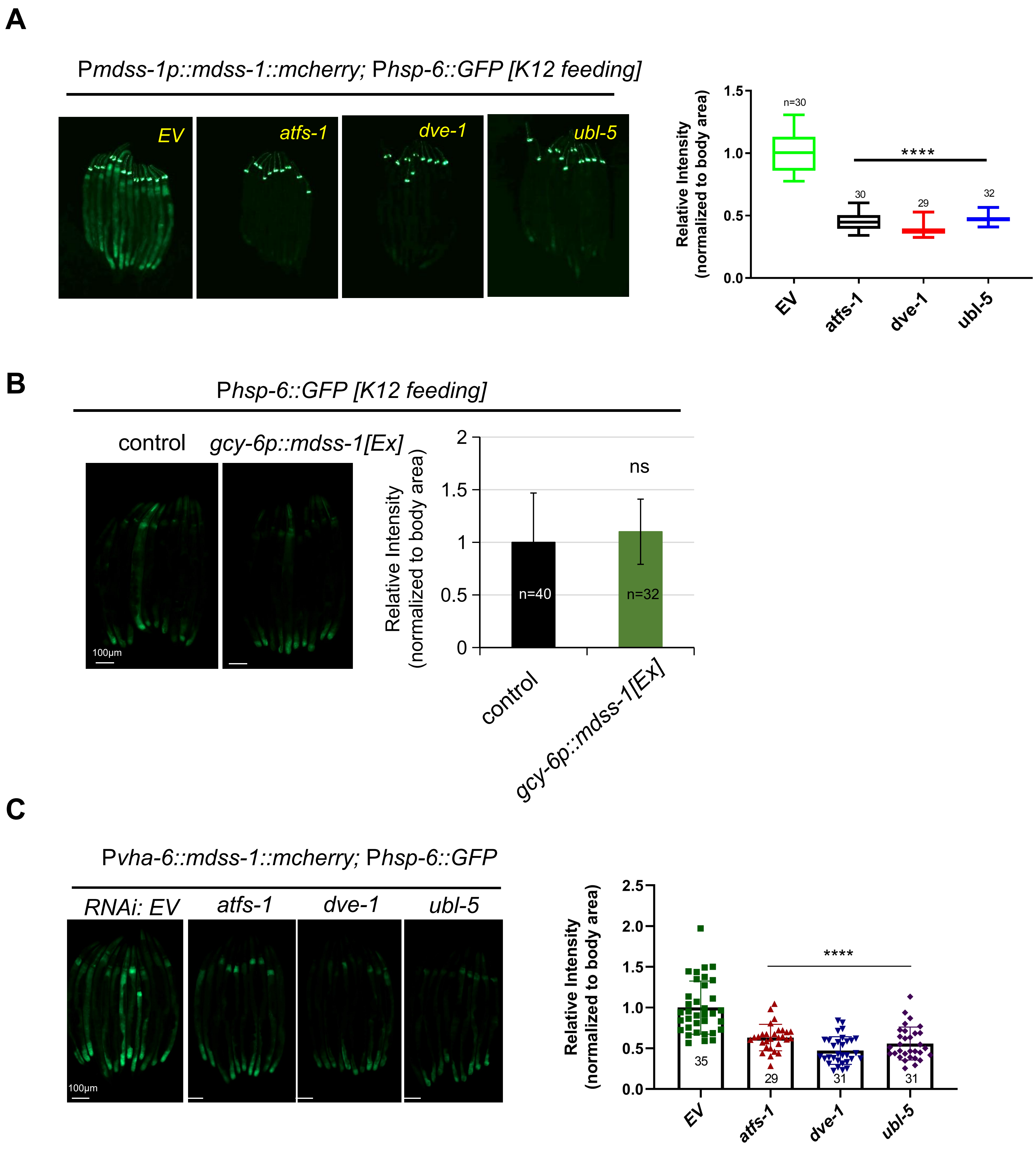


**Figure S5. MDSS-1 acts as a potential PGN receptor in $\text{UPR}^{\text{mt}}$ signal transmission. Relative to Figure 5.**

**(A)** Microscope images and fluorescence quantification of *Phsp-6::GFP* reporter expression in animals containing P*mdss-1::mdss-1::mcherry* grown on either empty vector (EV), *atfs-1* (RNAi), *dve-1* (RNAi) or *ubl-5* (RNAi). Animals carrying transgenes discriminated by P*odr-1::GFP.*

**(B)** Microscope images and fluorescence quantification of P*hsp-6p::GFP* reporter expression in animals with or without P*gcy-6::mdss-1[Ex]::mcherry* transgene grown on *E. coli* K12.

**(C)** Microscope images and fluorescence quantification of *Phsp-6::GFP* in animals carrying P*vha-6::mdss-1::mcherry* transgene grown on either empty vector (EV), *atfs-1* (RNAi), *dve-1* (RNAi) or *ubl-5* (RNAi).

For all panels, *p<0.05, **p<0.01, ***p<0.001, ****p<0.0001, n is the number of worms scored. Error bars, ± s.d. All experiments were performed independently at least three times.

Figure S6

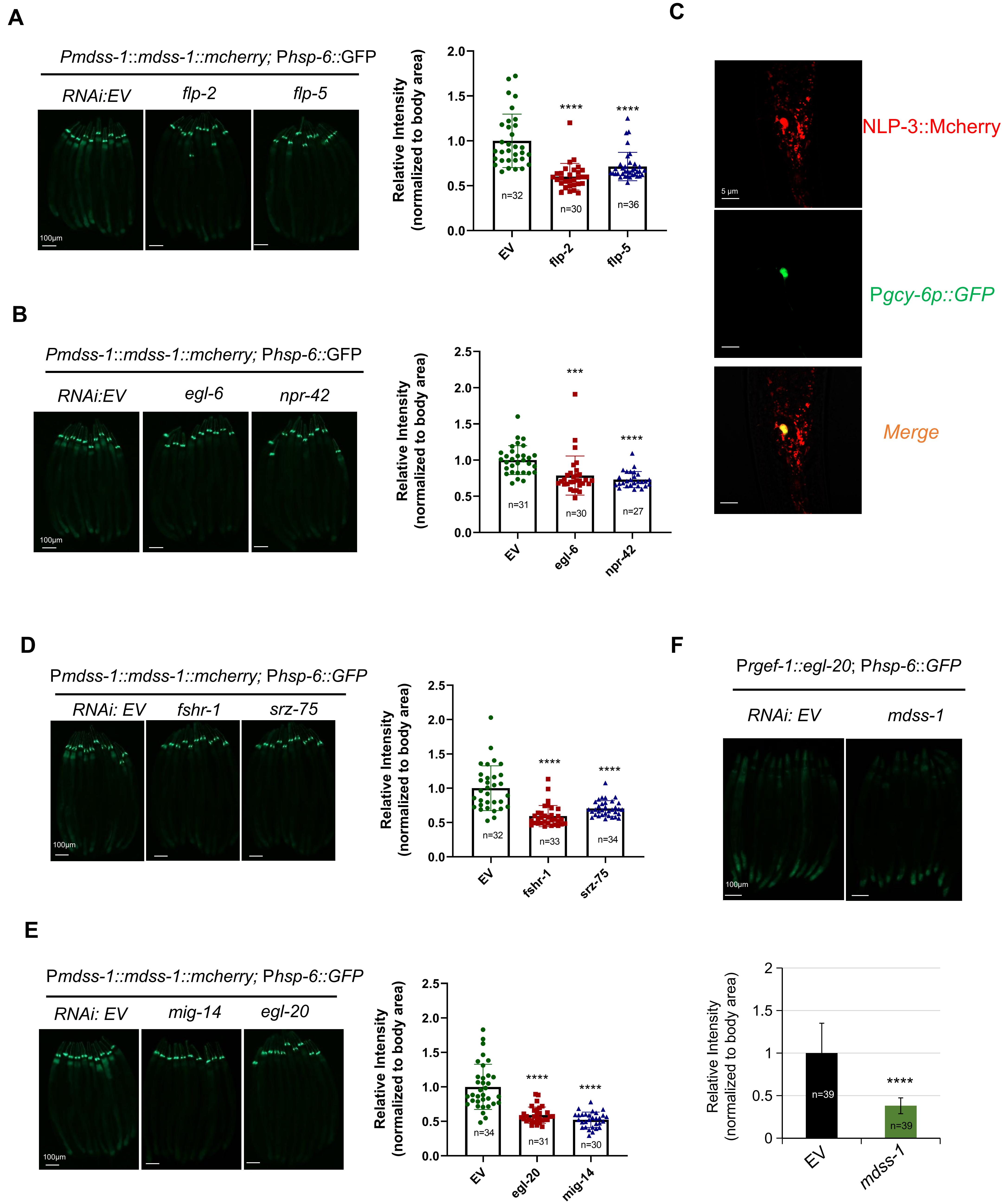
A
Pmdss-1::mdss-1::mcherry; Phsp-6::GFP
RNAi:EV
flp-2
flp-5
100μm
Relative Intensity (normalized to body area)
2.0
1.5
1.0
0.5
0.0
****
****
n=32
n=30
n=36
EV
flp-2
flp-5
B
Pmdss-1::mdss-1::mcherry; Phsp-6::GFP
RNAi:EV
egl-6
npr-42
100μm
Relative Intensity (normalized to body area)
2.5
2.0
1.5
1.0
0.5
0.0
***
****
n=31
n=30
n=27
EV
egl-6
npr-42
C
5 μm
NLP-3::Mcherry
Pgcy-6p::GFP
Merge
D
Pmdss-1::mdss-1::mcherry; Phsp-6::GFP
RNAi: EV
fshr-1
srz-75
100μm
Relative Intensity (normalized to body area)
2.5
2.0
1.5
1.0
0.5
0.0
****
****
n=32
n=33
n=34
EV
fshr-1
srz-75
F
Prgef-1::egl-20; Phsp-6::GFP
RNAi: EV
mdss-1
100μm
E
Pmdss-1::mdss-1::mcherry; Phsp-6::GFP
RNAi: EV
mig-14
egl-20
100μm
Relative Intensity (normalized to body area)
2.5
2.0
1.5
1.0
0.5
0.0
****
****
n=34
n=31
n=30
EV
egl-20
mig-14
Relative Intensity (normalized to body area)
2
1.5
1
0.5
0
****
n=39
n=39
EV
mdss-1

**Figure S6. The activation of MDSS-1-mediated UPR$^{mt}$ involves the participation of inter-tissue communication factors. Relative to Figure 6.**

**(A)** Microscope images and fluorescence quantification of P*hsp-6p::GFP* reporter expression in animals containing P*mdss-1::mdss-1::mcherry* transgene grown on empty vector (EV), *flp-2* (RNAi) or *flp-5* (RNAi). Animals carrying transgenes discriminated by P*odr-1::GFP*.

**(B)** Microscope images and fluorescence quantification of P*hsp-6p::GFP* reporter expression in animals containing P*mdss-1::mdss-1::mcherry* grown on empty vector (EV), *egl-6* (RNAi) or *npr-42* (RNAi). Animals carrying transgenes discriminated by P*odr-1::GFP*.

**(C)** Microscope images of wild-type animals carrying P*nlp-3::nlp-3::mcherry* and P*gcy-6::GFP*.

**(D)** Microscope images and fluorescence quantification of *Phsp-6p::GFP* reporter expression in animals containing P*mdss-1::mdss-1::mcherry* transgene grown on either empty vector (EV), *fshr-1* (RNAi) or *srz-75* (RNAi). Animals carrying transgenes discriminated by P*odr-1::GFP*.

**(E)** Microscope images and fluorescence quantification of P*hsp-6p::GFP* reporter expression in animals containing P*mdss-1::mdss-1::mcherry* transgene grown on either empty vector (EV), *mig-14* (RNAi) or *egl-20* (RNAi). Animals carrying transgenes discriminated by P*odr-1::GFP*.

**(F)** Microscope images and fluorescence quantification of P*hsp-6::GFP* reporter expression in animals containing *Prgef-1::egl-20* transgene grown on empty vector (EV) and *mdss-1*(RNAi).

For all panels, *p<0.05, **p<0.01, ***p<0.001, ****p<0.0001, n is the number of worms scored. Error bars, ± s.d. All experiments were performed independently at least three times.

### Figure S7

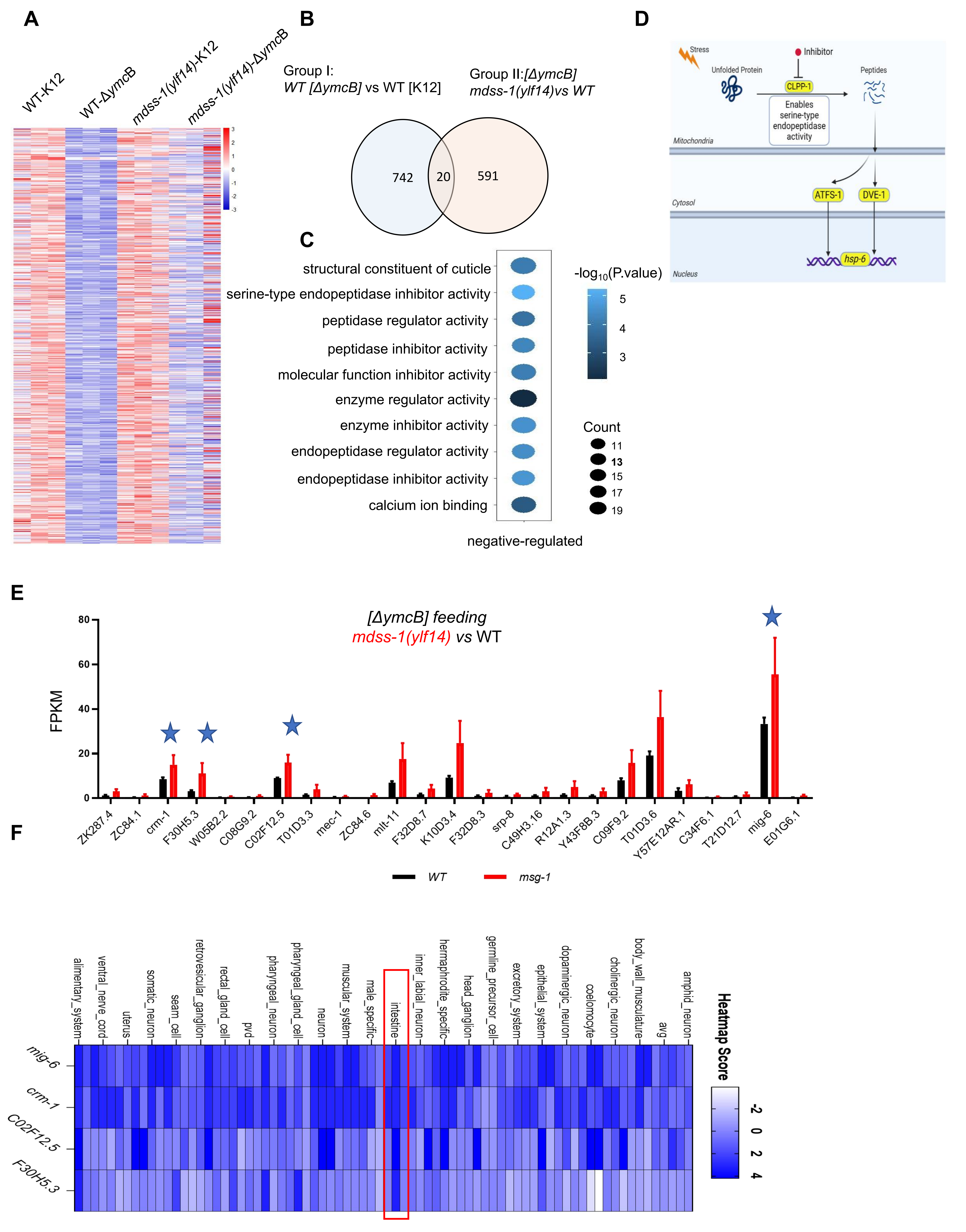
A
WT-K12
WT-ΔymcB
mdss-1(ylf14)-K12
mdss-1(ylf14)-ΔymcB
B
Group I:
WT [ΔymcB] vs WT [K12]
Group II:[ΔymcB]
mdss-1(ylf14)vs WT
742
20
591
C
structural constituent of cuticle
serine-type endopeptidase inhibitor activity
peptidase regulator activity
peptidase inhibitor activity
molecular function inhibitor activity
enzyme regulator activity
enzyme inhibitor activity
endopeptidase regulator activity
endopeptidase inhibitor activity
calcium ion binding
-log10(P.value)
Count
negative-regulated
D
Stress
Inhibitor
Unfolded Protein
CLPP-1
Peptides
Enables serine-type endopeptidase activity
Mitochondria
ATFS-1
DVE-1
Cytosol
hsp-6
Nucleus
E
[ΔymcB] feeding
mdss-1(ylf14) vs WT
FPKM
ZK287.4
ZC84.1
crm-1
F30H5.3
W05B2.2
C08G9.2
C02F12.5
T01D3.3
mec-1
ZC84.6
mlt-11
F32D8.7
K10D3.4
F32D8.3
srp-8
C49H3.16
R12A1.3
Y43F8B.3
C09F9.2
T01D3.6
Y57E12AR.1
C34F6.1
T21D12.7
mig-6
E01G6.1
WT
msg-1
F
alimentary_system
ventral_nerve_cord
uterus
somatic_neuron
seam_cell
retrovesicular_ganglion
rectal_gland_cell
pvd
pharyngeal_neuron
pharyngeal_gland_cell
neuron
muscular_system
male_specific
intestine
inner_labial_neuron
hermaphrodite_specific
head_ganglion
germline_precursor_cell
excretory_system
epithelial_system
dopaminergic_neuron
coelomocyte
cholinergic_neuron
body_wall_musculature
avg
amphid_neuron
mig-6
crm-1
C02F12.5
F30H5.3
Heatmap Score

**Figure S7. The activation of MDSS-1-mediated UPR$^{mt}$ involves the endopeptidase inhibitors. Relative to Figure 6.**

**(A)** Heatmap of deferentially down-regulated genes induced by *ΔymcB* in group I (wild-type animals grown on K12 followed by exposure to K12 or *ΔymcB* ) and group II (*mdss-1(ylf14)* animals grown on K12 followed by exposure to K12 or *ΔymcB*). Genes with an adjusted p value< 0.05 were selected as differentially expressed genes.

**(B)** Venn diagram of numbers of deferentially down-regulated genes in wild-type and *mdss-1(ylf14)* mutant animals grown on K12 transferred to *ΔymcB* for 24 hours. Genes with an adjusted p value< 0.05 were selected as deferentially expressed genes.

**(C)** GO enrichment of overlapped genes in Figure 7(D). Gene numbers are shown in circle size and P.values are shown in color. See also table S4.

**(D)** A schematic of CLPP-1 involved UPR$^{mt}$ in *C.elegans*. The activity of CLPP-1 that degrading the unfolded protein into peptide fragments was inhibited by serine-type endopeptidase inhibitors, which in turn affected the activation of UPR$^{mt}$ downstream.

**(E)** The genes with serine-type endopeptidase inhibitor activity, which are negatively regulated by MDSS-1, were identified through RNA-seq analysis from Group II.

**(F)** Heatmap Score of these endopeptidase inhibitors in different tissues. Low score is shown in white, and high score is in blue. See also Table S5.Data from worm tissue prediction application: https://worm.princeton.edu/ (Kaletsky R et al., 2018)
